# The evolution of parthenogen fertilization rates in switching environments: from facultative cell-fusion to oogamy

**DOI:** 10.1101/2023.01.13.523933

**Authors:** Xiaoyuan Liu, Jon Pitchford, George W.A. Constable

## Abstract

Organisms with external fertilisation exhibit a broad range of reproductive modes, from simple parthenogenesis to sexual reproduction encompassing isogamy, anisogamy, and oogamy, and including environmentally-mediated facultative sex. Here we develop a unifying mathematical model which explains the emergence of these modes via the coevolution of fertilization rate and cell size. Using a minimal assumption that survival is dependent on cell mass, and by carefully accounting for biological and evolutionary time scales, we find two distinct evolutionary outcomes: high fertilization rate (obligate sexuality) is selected when costs to cell fusion are low, while zero fertilization rate (obligate asexuality) is selected for when these costs are high. Surprisingy, in high fertilization rate scenarios evolving populations can transition from isogamy to anisogamy and oogamy via evolutionary branching. Furthermore, in variable environments we show that, without phenotypic plasticity, intermediate fertilization rates and isogamy can be maintained through bet-hedging. Allowing phenotypic plasticity can give rise to facultative sex; sexual reproduction in harsh environmental conditions, and asexuality in more benign conditions. These results parsimoniously explain a large range of empirically observed parthenogen reproduction strategies, and offer an hypothesis for the origin of binary cell fusion, a key step in the evolution of syngamy and sexual reproduction itself.

## 1. Introduction

Fertilization is a crucial step in reproductive cycle of all sexually reproducing organisms. The modelling of fertilization kinetics is an interesting topic in its own right [1, 2], and has been argued to be a key component in the evolution of anisogamy (unequal gamete sizes) from isogamy (equal gamete sizes) [3, 4]. Traditionally, models assume obligate sexual reproduction (unfertilized gametes die at the end of each generation) and a fixed fertilization rate (the rate at which gametes encounter one another and fuse to form zygotes) [5, 6, 7], with the further evolution of oogamy (motile microgametes and sessile macrogametes) possible when factors such as speed-mass relationships [8], costly motility [9] or internal fertilization are accounted for [10]. Recently, however, inspired by the life-histories of green and brown algae such as *Blidingia minima* (isogamous [11]), *Urospora neglecta* (anisogamous [12]), and *Saccharina japonica* (oogamous [13]), whose gametes can develop parthenogenically should they fail to find a mate (see [11, 12, 14], respectively), the first of these assumptions was relaxed in two theoretical papers [15, 16]. In [15], extra survival costs were incurred by gametes developing parthenogenically, while in [16] extra survival costs on either the parthenogenic or the sexual reproductive route were considered. These studies lead to a natural question; if the sexual route (via fertilization) and asexual route (via parthenogenesis) carry different survival costs, how should the fertilization rate evolve to account for this? Should this rate increase (to minimise the number of unfertilized gametes taking a potentially perilous route to survival) or should it decrease (to avoid potential costs incurred during cell fusion)? Here we tackle this question, and arrive at two potential evolutionary outcomes; selection for high fusion rates (tending to obligate sexuality) when costs to cell fusion are low to moderate, and selection for zero fusion rate (functionally obligate asexuality) when costs to cell fusion are high.

These results inform the question of the evolution of sexual reproduction itself, where the majority of theoretical work has focused on the benefits and costs of genetic recombination. The benefits of recombination arise from generating individuals with beneficial gene combinations that would not have arisen by mutation alone [17]. However, this process simultaneously incurs a cost; a sexual population breaks down its beneficial gene associations more than would an asexual population [18]. Further, the act of sex is time and energy intensive and necessitates locating a partner [19]. In these theoretical studies, the costs of recombination often outweigh the benefits, in contradiction to the empirically observed success of sexual reproduction [20, 21, 22, 23, 24]. Here we do not account for such genetic factors. Rather, we note that although the details of the early evolution of sexual reproduction in the last common eukaryotic common ancestor (LECA) are shrouded in mystery, it is argued that the process began with the evolution of cell–cell fusion and meiosis [25]. This step can be further broken down into the evolution of binary cell fusion, the one spindle apparatus, homologous pairing and chiasma, and finally reduction, division and syngamy [26].

Traditional hypotheses for the evolution of binary cell fusion generally rely on hybrid fitness advantage. It has been suggested that selection for cell–cell fusions might have initially been driven by “selfish” transposons and plasmids [27, 28, 29], or negative epistatic interactions between mitochondrial mutations [30, 31]. However, once a heterokaryotic cell has been formed (binucleate with nuclei from both parental cells), the advantage of hybrid vigor and the masking of deleterious mutations could lead to the maintenance of cell fusion [26]. Such benefits are required to alleviate costs to cell-fusion, which include selfish extra-genomic elements in the cytoplasm [32] and cytoplasmic conflict [33, 34].

In contrast, the hypothesis that we put forward here, namely that the survival advantage that comes from increasing cell-size might select for binary cell fusion, relies on the physiological advantages conferred by cell-cell fusion and is independent of the question of the genetic advantages (and disadvantages) of sexual reproduction. This benefit of increasing cytoplasmic volume has been explicitly identified in the landmark paper introducing the Parker, Baker and Smith (PBS) model for the evolution of anisogmay [3], but has not yet been used to investigate the evolution of binary cell fusion itself. In providing a mechanistic hypothesis for the evolution of binary cell fusion, our work echoes [35], where an advantage to cell fusion is identified in terms of shortening the cell-cycle.

Sexual reproduction is correlated with challenging environmental conditions in many faculatively sexual organisms [36, 37]; for instance sexual reproduction is triggered by nitrogen starvation in the green algae *Chlamydomonas reinhardtii* [38] and yeast *Saccharomyces pombe* [39], and darkness, moisture and phosphate starvation in the case of the social amoeba *Dictyostelium discoideum* [40]. In extending our model to account for changing environments, our work follows many theoretical papers on the evolution of sexual reproduction [41, 42, 43, 44] and mating types [45]. Because the evolving traits in our model (cell size and fertilization rate) are continuous, we employ techniques from adaptive dynamics [46] generalised to dynamic environments [47].

The paper is organised as follows. In Section 2 we introduce the model. The key assumptions are described in Section 2.1, which concludes with the derivation of the evolutionary dynamics of a population in a fixed environment. In Section 2.2 the evolutionary dynamics are modified to account for switching environments, and the role of phenotypic plasticity in this context is evaluated in Section 2.3. The evolutionary dynamics are analysed in Section 3. We conclude by discussing the biological interpretation of these results in Section 4, along with discussing potential future work.

## 2. Model

In this section we describe the specifics of the models we use, paying specific attention to the various time scales involved. We begin by considering the evolutionary dynamics in a fixed environment (Section 2.1), before generalising to the case of a fixed phenotype in a dynamic environment under which bet-hedging strategies can evolve (Section 2.2). We finally consider the case of a plastic phenotype in a dynamic environment (Section 2.2).

### 2.1. Model dynamics in a fixed environment

The evolutionary dynamics of the model are built from a hierarchy of timescales, which are particularly important to keep in mind once environmental switching is introduced in later sections. The shortest timescale is the generational timescale. The intermediate timescale is that over which the invasion of a rare mutant (taking place over many generations) can occur. The longest timescale is the evolutionary timescale, representing the cumulative effect of multiple mutations and invasions.

#### Dynamics within each generation

At the start of each discrete generation, a number o: adults (mass M) produce daughter cells (mass *m*), such that mass is conserved (i.e. each adult produces *M/m* daughter cells). Note that this implicitly assumes, for simplicity, that a continuous (rather than discrete) number of daughter cells is possible. Daughter cells then enter a pool and the fusion (fertilization) process takes place. After a finite time window, the resultant cells face a round of survival dependent on their mass. The surviving cells form the basis of the next generation, completing the generational cycle, as illustrated in Figure 1.

**Figure 1:**
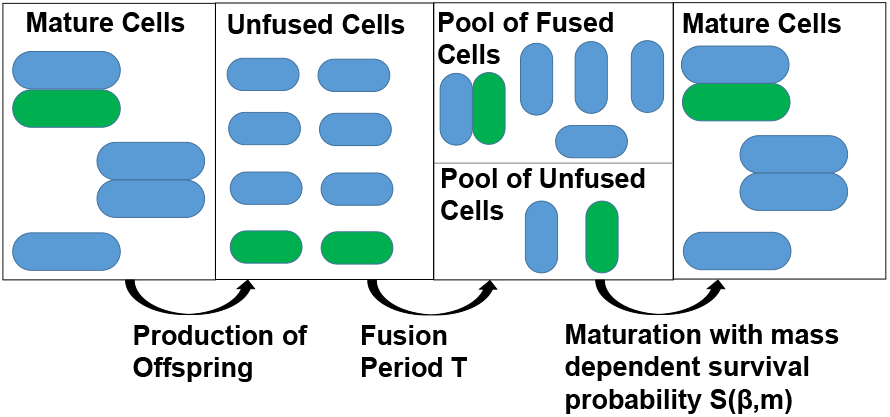
Schematic of dynamics within each generation. Mature cells produce daughter cells at the start of a generation. All the daughter cells are given a fixed time period *T* in which to complete the fusion process. At the end of the fusion period, there will be a pool of fused and unfused cells, both of which are capable of maturation. Each cell survives according to its independent survival functions (*S_z_*(*β*) and *S_p_*(*β*) respectively) to produce a number of mature cells in the subsequent generation. The pool of unfused daughter cells consist of (blue) and mutant gametes (green), where the mutation occurs in either the mass *m* or fusion rate *α*.

The model can be usefully interpreted in two distinct ways, both of which are relevant for the biological consequences of the results:

##### Eukaryotic parthenogens

Adults produce haploid self-compatible gametes (i.e. we do not model self-incompatible mating types). These gametes are placed in a pool and undergo a period of fertilisation according to mass-action dynamics. Fertilised gametes produce diploid zygotes, while unfertilised gametes can develop as haploids parthenogenically (e.g. parthenosporophytes [48]). The survival of diploid zygotes and haploid parthenosporophytes is dependent on their mass. Following survival, one may view the zygotes as undergoing meiosis to produce haploid adults (e.g. gametophytes), or developing as diploid adults (e.g. sporophytes) for the next generation [49]; assuming no fitness differences arising from ploidy, these interpretstions are mathematically equivalent up to a generational delay [16]. Adults then produce haploid gametes, completing the generational cycle.

##### Evolution of binary cell fusion

Adults produce a number of daughter cells, each containing a single haploid nucleus. These daughter cells are placed in a pool and undergo a period in which they may fuse to form a binucleated cell (e.g. a dikaryon, in which the cytoplasms of the contributing cells are mixed but their nuclei remain distinct [50]) or remain a mononucleated cell. The survival of these two cell-types is a function of their mass, plus an additional survival risk to binucleated cells. Surviving adults divide to form a new generation of mononucleated haploid daughter cells, with binucleated parental cells producing mononucleated progeny through vegetative segregation [51]; note that although we do not account for the possibility of binucleated cells failing to form mononucleated progeny, this can be accounted for by their additional survival cost.

In light of this generality, we refer to the cells produced by adults as daughter cells (rather than gametes) and the rate at which cells fuse as the fusion rate (rather than fertilization rate). We describe these processes in more mathematical detail below.

##### Fusion Kinetics

We assume that all cells may fuse with each other (i.e. there are no self-incompatiable mating types). Note that this assumption is consistent with most models of the early evolution of sexual reproduction, which suppose the existence of a “unisexual” early ancestor that mated indiscriminately [52]. Initially, the population comprises *N* single daughter cells. We assume fertilisation is external, with cells fusing according to mass action dynamics at a fusion rate *α*, such that the number of single (unfused) cells, *N*, is given by the solution to

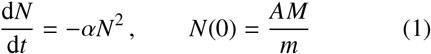

(see also [53]). At the end of the fusion window or duration *T* there are then *N*(*T*) single cells remaining, and (*N*(0) − *N*(*T*))/2 fused cells.

The parameter *α* is variously referred to as the fertilization rate, the collision rate [1], the “‘aptitude’ for union” [54], or the “bimolecular reaction constant” [55]. We will refer to *α* as the fusion rate, and treat it as a trait subject to evolution. Overall, *α* captures the compound effect of the propensity for fusion between cells encountering each other, as well as additional mechanisms to enhance cell encounter rate such as increased emission or response to cheomoattractants [56] or cell motility [57]. In practice there is a likely upper-bound on *α*, brought about by energy trade-offs, diffusion in aquatic environments, or dispersal in terrestrial environments. To account for this we introduce a ceiling on *α*, and restrict its evolutionary dynamics to the range *α*_max_ ≥ *α* ≥ 0.

##### Survival Probability

At the end of a finite fusion window at time *T*, the population will consist of both fused and unfused cells. We assume that the probability that a either of these cell types survives is given by the Vance survival function [58], which is a common assumption in the literature [59, 5, 7]. Given a cell size *m_c_* (for either fused or unfused cells), the survival probability is given by

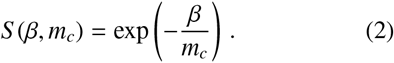

Note that this is an increasing function of cell size, and we do not account for daughter-cell mortality during the fusion period. Thus, although both fused and unfused cells are exposed to the same survival function, fused cells (with a mass around twice the size of unfused cells in a monomorphic population), will have a greater survival probability than unfused cells. Meanwhile for a given mass *m_c_*, increasing *β* will decrease the survival probability. We therefore refer to *β* as the resistance to survival, with high *β* corresponding to harsh environments in which survival is difficult, and low *β* corresponding to more benign environments in which even cells of modest mass have a high probability of surviving.

In addition to the benefits explained in the introduction, cell fusion may also carry risks: generally these include cell-fusion failure [60], selfish extra-genomic elements in the cytoplasm [32] and cytoplasmic conflict [33, 34]; in eukaryotes sexual reproduction carries both genetic and energetic costs [61]; in the evolution of early binary cell fusion, costs could include the maintenance of a binucleated cell [62] and the cost of failed segregation. We therefore introduce an additional fixed cost, 1 ≥ *C* ≥ 0, applied to fused cells independent of their mass. The final probability of survival for a cell formed from the fusion of two daughter cells of size *m* is then given by (1 – *C*) exp [–*β*/(2*m*), while the probability of survival for an unfused cell of size *m* is exp [–*β*/*mt*].

#### Invasion Dynamics

We assume that haploid daughter cells are characterised by two genetically determined non-recombining traits; their mass, *m*, and their fusion rate, *α*. We next consider a monomorphic resident population to which a mutant individual is introduced at rate *μ.* This mutant may produce daughter cells of a different mass to its ancestor, *m* ± *δm*, where *δm* represents the size of a mutational step. Under this scenario the mutant may produce more or fewer daughter cells than its ancestor (see Appendix A.1), but the survival probability of its unfused cells (mass *m* ± *δm*), mutant-resident fused cells (mass 2*m* ± *δm*), and mutant-mutant fused cells (mass 2(*m* ± *δm*)), will also be simultaneously decreased or increased (see Eq. (2) and Appendix A.3). The cumulative effect of this quality-quantity trade-off will either lead to selection for or against the mutant over subsequent generations.

Alternatively, the mutant may engage in an increased or decreased fusion rate relative to its ancestor, at a rate *α* ± *δα* (see Appendix A.2). Under this scenario the mutant fuses with residents at their average fusion rate (2*α* ± *δα*)/2 and other mutants at a rate *α* ± *δα*. Mutants will either contribute to more or fewer fused cells, and depending on the resistance to survival, *β*, and the cost to cell fusion, *C*, may experience a selective advantage over the resident by devoting more of its daughter cells to one of the reproductive routes (see Appendix A.4).

In order to mathematically characterise the invasion dynamics (which occur over discrete generations), we assume that *δm* and *δα* are small, so that we can approximate the dynamics continuously. Denoting by 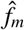 the frequency of mutants of size *m* ± δ*m* in the population, and *t_g_* the number of generations, we find (see Appendix B.1)

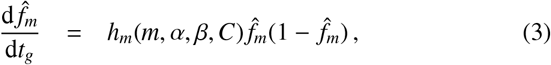

where *h_m_*(*m, α,β, C*) is a constant that depends on the parameters *m,α,β, C* (see Eq. (B.1)). Similarly, denoting by *f_α_* the frequency of mutants with fusion rate *α* ± *δα* in the population (see Appendix B.2), we find

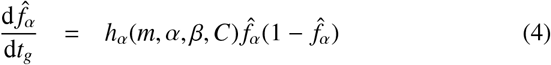

where *h_α_*(*m, α,β, C*) is a constant that depends on the parameters *m, α,β, C* (see Eq. (B.2)). We thus see that in the case of a single mutant, we have frequency-independent selection for both traits, with the direction of selection determined by the resistance to survival, *β*, the cost to cell fusion, *C*, and the traits of the resident population, *m* and *α*. We assume for the remainder of the mathematical analysis that mutants encounter a strictly monomorphic resident population, allowing us infer a simple trait substitution process which significantly simplifies the subsequent evolutionary analysis. (Simulations accounting simultaneously for many genetic variants provide important new insight, see Appendix C.3 and discussion below).

#### Evolutionary Dynamics

Following standard approaches in adaptive dynamics (see also Appendix C), we construct the evolutionary equations for the daughter cell size, *m*, and the fusion rate, *α*. Denoting by *τ* the evolutionary timescale over which mutations appear and trait substitutions occur, we find

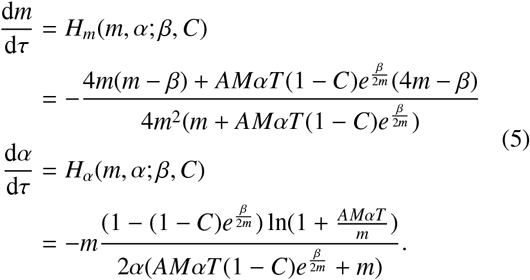

Note that these are highly nonlinear functions, with some evident non-physical behaviour near the boundaries (e.g. when cell masses are zero). This feature is common in simplified models of anisogamy evolution [16]; we comment on these further in the Results, Section 3.

### 2.2. Evolutionary dynamics in switching environments without phenotypic plasticity

We now wish to consider the case of a population characterised by daughter cell mass (*m*) and fusion rate (*α*) traits, evolving subject to changing environmental conditions. Explicitly, we allow the resistance to survival to alternate between two values *β*_1_ and *β*_2_; recall from Eq. (2) that if *β*_1_ > *β*_2_ then *β*_1_ represents a comparatively “harsh environment”, where cells have a lower survival probability than in environment *β*_2_.

Switching between these two environments is modelled as a discrete stochastic telegraph process [63, 45]; the time spent in each environment is distributed geometrically (a discrete analogue of the exponential distribution), spending an average period *τ*_1_ ≈ 1/λ_1→2_ in environment 1 and τ_2_ ≈ 1/λ_2→1_ in environment 2, where *λ*_*i*→*j*_ is the transition rate from environment *i* to *j*. We must carefully consider the magnitude of these timescales in comparison with the other timescales at work in the model (see Section 2.1).

First consider the case where the environmental switching timescales, *τ*_1_ and *τ*_2_, are larger than the generational timescale (*t_g_*), but much smaller than the invasion timescale (characterised by the inverse of the strength of selection, proportional to 1/*δ_m_* and 1/*δ_α_*) and the mutational timescale (1/*μ*). We call this the “fast relative to invasion” switching regime (FRTI). In this scenario, the population does not switch environments during a single round of fusion kinetics, but typically switches between the two environmental states many times before an invasion has time to complete. When switching occurs this frequently, we can approximate the dynamics mathematically by observing that the population experiences the weighted average of the selection pressures in the two environments [63]. Denoting by *P*_1_ and *P*_2_ the probability of finding the population in either of the respective environments, we have

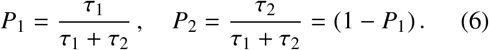

The effective selection pressure on mutants with mass *m* + *δm* in a resident population of mass *m* during an invasion is then given by

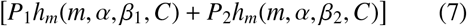

which can be contrasted against the selection pressure *h_m_*(*m, α, β, C*) in Eq. (3). An analogous approach allows us to approximate the invasion dynamics for mutants with a different fusion rate to their ancestors in this FRTI regime (see Appendix D.1).

With equations for 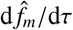 and 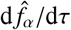 in hand, we can proceed to apply the same standard techniques from adaptive dynamics as were used to derive Eq. (5) from Eqs. (3-4) (see Appendix D). We obtain the effective evolutionary dynamics

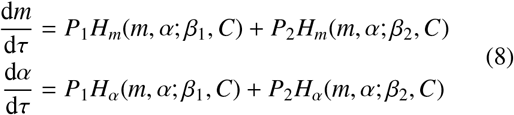

where *H_m_*(*m, α*; *β, C*) and *H_α_*(*m, α*; *β*_1_, *C*) retain the functional forms given in Eq. (5), and *P*_1_ and *P*_2_ are taken from Eq. (6).

In contrast to the FRTI regime discussed above, we can also investigate the regime in which environmental switching occurs on a comparable or slower timescale than invasion, but still occurs fast relative to the evolutionary timescale. In this scenario, which we call the “fast relative to evolution” switching regime (FRTE), environmental switching occurs on a similar rate to that at which new mutations are introduced, but much faster than the combined effect of mutation and selection (e.g. *λ_i→j_* ≪ *μ* × *δ_m_*), such that only a small number of mutations can fixate in either environment before the population switches to the alternate environment. Although we shall show via simulations that this FRTE regime leads to quantitatively different evolutionary trajectories compared to the FRTI regime, we show mathematically in Appendix D.2 that the evolutionary dynamics can be approximated by the same equations (see Eq. (8)).

### 2.3. Evolutionary dynamics in switching environments with phenotypic plasticity

We now consider environmental switching implemented in an identical manner to the previous Section 2.2, but we allow the population to evolve different strategies in response to the two environments. Accounting for this phenotypic plasticity, the population’s evolutionary state is now described by four traits; the daughter cell mass in environments 1 and 2 (*m*_1_ and *m*_2_) and fusion rate in these environments (*α*_1_ and *α*_2_).

For simplicity we assume that any cost of phenotypic switching or environmental sensing is negligible and that this plastic switching is instantaneous upon detection of the change in environmental conditions. We show in Appendix E that the dynamics are then given by

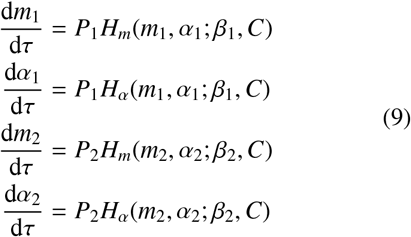

where *H_m_*(*m,α*;*β, C*) and *H_α_*(*m,α*;*β*_1_, *C*) again retain the functional forms given in Eq. (5). Finally, we assume that the initial evolutionary dynamics must begin from a phenotypically undifferentiated state in which *m*_1_(0) = *m*_2_(0) ≡ *m*(0) and *α*_1_(0) = *α*_2_(0) ≡ *α*(0).

## 3. Results

In this section we proceed to analyse the evolutionary dynamics derived in Sections 2.1-2.3 and compare our results to numerical simulations of the full stochastic simulations.

### 3.1. In a fixed environment the population evolves to either fully sexual or fully asexual reproduction

In Figure 2, we see two potential evolutionary outcomes for the co-evolutionary dynamics of *m* and *α* in a single fixed environment that are dependent on the initial conditions and parameters; the population can either evolve to large (technically infinite) fusion rates or to zero fusion rates.

**Figure 2:**
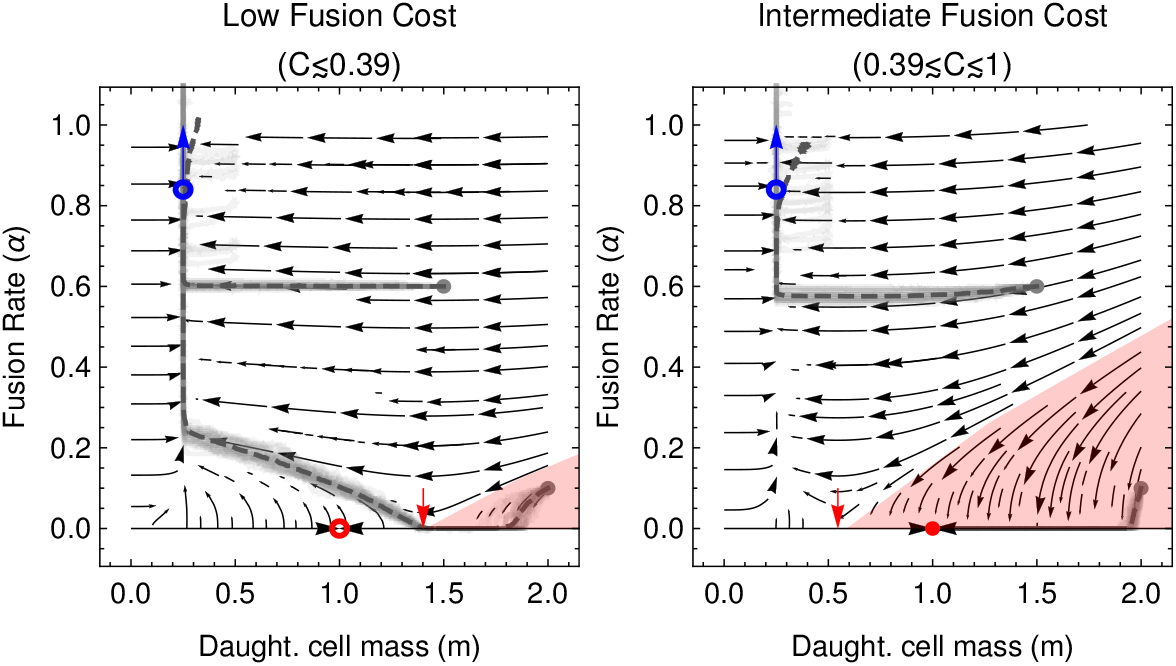
Phase portraits for the co-evolutionary dynamics in a fixed environment (see Eq. (5)). High fusion rates (obligate sex) are the only evolutionary outcome when costs to cell fusion are low (panel (a)), while under high costs, high fusion rate and zero fusion rate (obligate asex) are both evolutionary outcomes, as summarised analytically in Eq. (10). The red shaded region shows trajectories leading to points on the *α* = 0 boundary for which evolution selects for decreasing fusion rate (d*α*/d*τ* < 0), while the critical point on this boundary at which d*α*/d*τ* = 0 is marked by the red arrow (see Appendix E.2). Red circles mark a fixed point in the evolutionary dynamics of *m* (*m*\* = *β*, see Eq. (C.2)), which may be unstable (open circles) or stable (filled circle) under coevolution with *α*. The blue circles and arrows illustrate the evolutionary attractor for high fusion rates ((*m*,α*\*) → (*β*/4, ∞), see Eq. (C.4)). Average population trait trajectories, (〈*m*〉(*t*), 〈*α*〉(*t*)), from simulation of the full stochastic model (see Appendix Appendix C.3) are plotted in light gray, and their mean over multiple realisations are given dark gray. The initial points of the trajectories are (*m*(0), *α*(0)) = (1.5,0.6) and (*m*(0), *α*(0)) = (2,0.1). The cost to fusion is *C* = 0.3 (panel (a)) and *C* = 0.6 (panel (b)). Remaining model parameters are *A* = 100, *M* = 1, *T* = 1 and *β* = 1. Simulation parameters are *δ* = 0.005, *f*_0_ = 0.002, *μ* = 1/2000, run for 5500/*μ* generations in (a), 6200/*μ* generatins in (b) and 3500/*μ* in (c).

When costs to fusion, *C*, are low (Figure 2, Panel a), there exists a smaller region of initial conditions that drive *α* towards zero (pink shaded region). When *α* = 0 within this region, selection on daughter cell-size, *m*, drives the population towards the point *m* = *β* (red dot, see also Appendix C.2). As this point exists outside the region in which d*α*/d*τ* < 0, selection for increased α can again manifest along the evolutionary trajectory. Thus when costs are sufficiently low, high fusion rates are the only evolutionary outcome.

Conversely when costs to fusion, *C*, are intermediate (Figure 2, Panel b), there exists a larger region of initial conditions that drive *α* towards zero (pink shaded region). The point *m* = *β* (red dot), towards which the population evolves when *α* = 0, is now contained within this region in which d*α*/d*τ* < 0, and so is a stable fixed point. Thus when costs are sufficiently high, there are two evolutionary outcomes, depending on the initial conditions; either high fusion rates or zero fusion rates.

Finally when costs to fusion, *C*, are extremely high, selection for decreased fusion rate acts regardless of initial conditions and a state in which *α* = 0 (zero fusion rate) is the only evolutionary outcome.

In Appendix C.2 we conduct a mathematical and numerical analysis to formalise the arguments above. In summary, the possible early evolutionary attractors, (*m*\*, *α*\*), are given by:

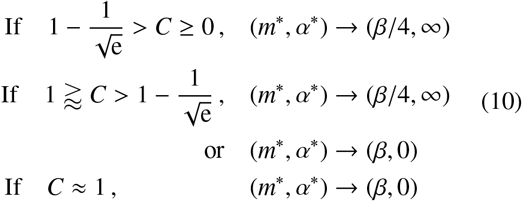

where we note 1 – e^-1/2^ ≈ 0.39. While intermediate costs 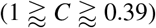 lead to two potential evolutionary outcomes depending on the initial conditions, it is the second of these, (*m*\*,*α*\*) = (*β*, 0), that is arguably the most relevant for the evolution of early cell fusion; if evolution had acted on daughter cell size, *m*, before the physiological machinery necessary for cell fusion had evolved, the initial condition for the co-evolutionary dynamics would be (*m*^0^, *α*^0^) → (*β*, 0), at which the population would be subsequently held by costs to fusion.

We conclude this section by addressing the key biological result that arises from this analysis; cell fusion is uniformly selected for even under moderately high costs (with a fraction of up to *C* ≈ 0.39 of fused cells failing to survive) and can even be selected for under higher costs (up to *C* ≈ 1) given necessary initial conditions. In the context of the evolution of early binary cell fusion, this provides a surprising nascent advantage to cell fusion. This advantage could even help compensate for other short-term costs arising from the later evolution of sex and recombination. The selective advantage experienced by fusing cells comes from their increased cytoplasmic volume, which leads to increased survival probabilities. Note that although we have not accounted for any costs for parthenogenic development [64], adding such costs would simply accentuate the observed behaviour.

### 3.2. At high fusion rates, the population can evolve from isogamy to anisogamy and finally oogamy

In Figure 2 we also see that the approximation obtained for the co-evolutionary dynamics of *m* and *α* in a single fixed environment, Eq. (5), accurately captures the dynamics of the full model realised via numerical simulation at early times. One point of departure is that at long times as *α* increases towards the (*β*/4, ∞) attractor predicted analytically, we see the mean mass trait value from simulations increasing to higher values than those predicted analytically. In Figure 3, we show that this is a result of evolutionary branching of the mass trait in the simulations. While we do not account for this behaviour in our analysis, it is consistent with the analytic criteria that trait selection should tend to zero at a branching point [65]; on inspection of *H_α_* (*m*, *α*; *β, C*) in Eq. (5), we see that d*α*/d*τ* → 0 with increasing *α*. The smaller “microgametes” evolve towards a mass *m*_micro_ = *δm* (the smallest mass allowed in the simulations) while the larger “macrogametes” evolve towards a mass *m*_macro_ = *β*. The initial branching described above can be biologically interpreted as selection for anisogamy, has been noted in other models that do not account for self-incompatible mating types [66].

**Figure 3:**
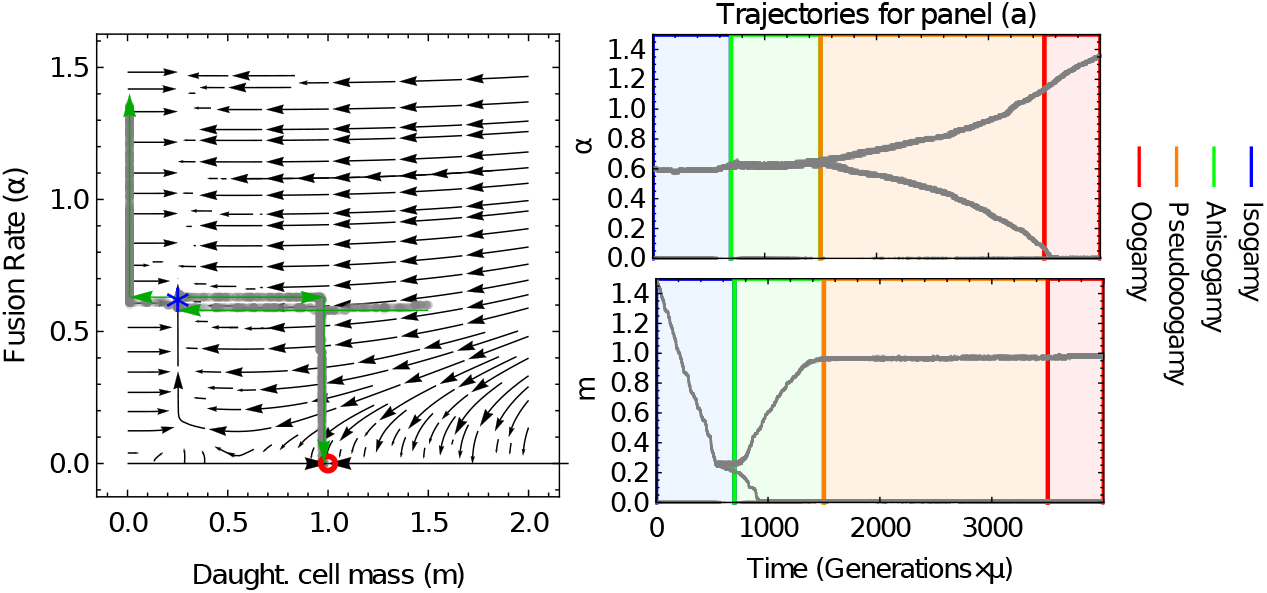
Numerical illustration of evolutionary branching in 2(b). Panel (a): Analytic predictions for the early evolutionary dynamics (as 2(b)) overlaid with trajectories (*m_i_*(*t*), *α_i_*(*t*))) for each *i*^th^ trait. Evolutionary branching is observed along the *m* = *β*/4 manifold, indicated by the blue star. Green arrows show the temporal progression of the branching. Panels (b) and (c): The temporal trajectories of the traits *α_i_*(*t*) and *m_i_*(*t*) respectively, showing that the evolutionary trajectory passes from isogamy to oogamy. Parameters used are *A* = 100, *M* = 1, *T* = 1, *C* = 0.6, *β* = 1, *δ* = 0.01, *μ* = 1/1000, *f*_0_ = 0.002, simulation run for 3500/*μ* generations.

**Figure 4:**
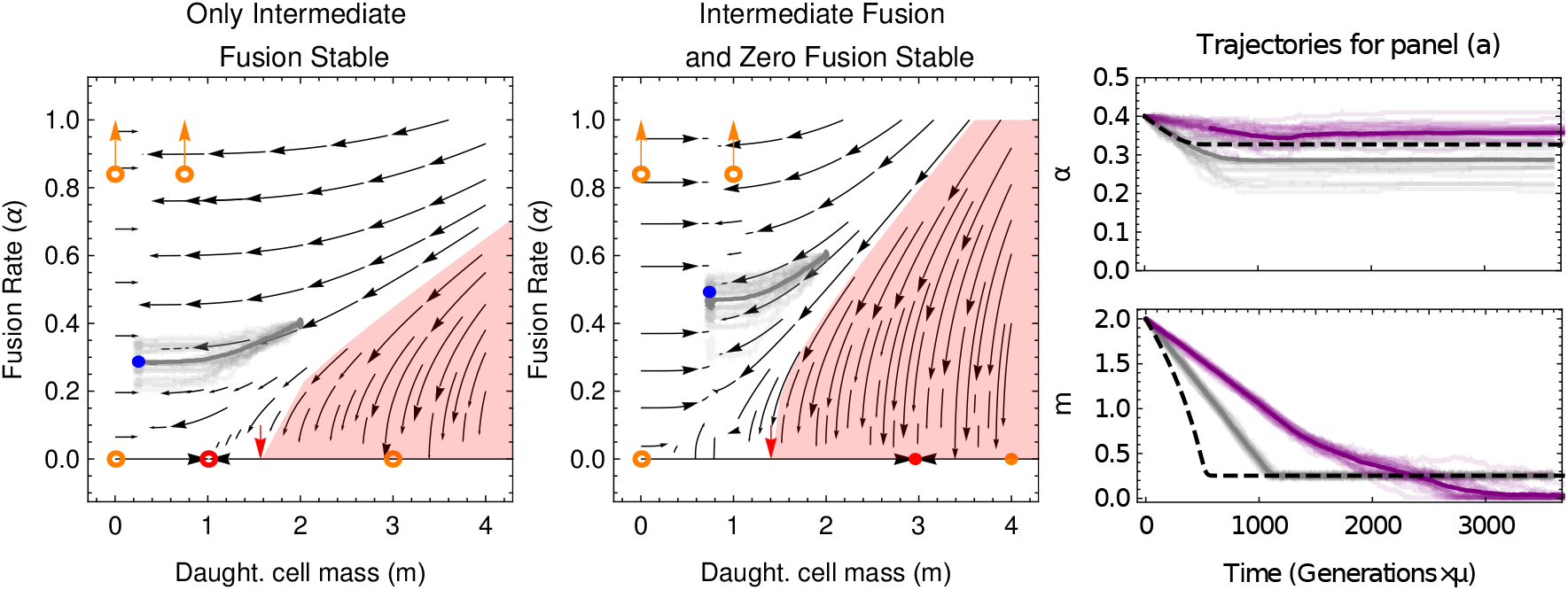
Phase portraits for the co-evolutionary dynamics in a switching environment without phenotypic switching (see Eq. (8)). In addition to qualitatively similar dynamics as in the fixed environment (see Figure 2), two new evolutionary scenarios are now possible, including populations in which stable intermediate fusion rates (filled blue circle) are the only evolutionarily stable state (see panel (a)) and populations in which there is an additional zero fusion stable state (filled red circle, see panel (b)). Open orange circles represent the (now unstable) states to which the population can be attracted in either environment 1 or two (where *β* = *β*_1_ or *β* = *β*_2_). Results of simulation of the full simulation model in the FRTI regime are overlaid in gray. In panel (c) we plot the time-series in the FRTI regime (gray) alongside those in the FRTE regime (purple) and our analytic predictions (black-dashed). In all panels *A* = 100, *M* = 10, and *T* = 0.1. In panel (a), *C* = 0.35, *β*_1_ = 3 (harsh environment), *β*_2_ = 0.01 (benign environment), (*m*(0), *α*(0)) = (2,0.4), and *P*_1_ = 0.335. In panels (b) and (c), *C* = 0.7, *β*_1_ = 4 (harsh environment), *β*_2_ = 0.01 (benign environment), (*m*(0), *α*(0)) = (2, 0.6), and *P*_1_ = 0.74. For the FRTI case, the mutation rate is *μ* = 0.00035 and for the FRTE case, *μ* = 0.0005. Stepsizes are δ = 0.007 for the FRTI case and *δ* = 0.005 for the FRTI case. In panel (a), *λ_GB_* = 67/532, *λ_BG_* = 1/4 for the FRTI case and *λ_GB_* = 67/1064000, *λ_BG_* = 1/8000 for the FRTE case. In panel (b), *λ_GB_* = 1/6, *λ_BG_* = 13/222 for the FRTI case and *λ_GB_* = 1/12000, *λ_BG_* = 13/444000 for the FRTE case. In all cases, the initial frequency of an invading strain is *f*_0_ = 0.002.

As the mass of the macrogamete becomes larger, a secondary branching in fusion rates occurs (see Figure 3(b)). Near this mass *m*_macro_ = *β*, the macrogamete has a high survival rate under parthenogenesis, and the benefits of cell fusion (in particular with microgametes of very small mass) are outweighed by the costs of cell fusion. Selection thus acts to lower the fusion rate of macrogametes, amacro, towards zero. This leads to an increased selection pressure for the microgametes to increase their fusion rate, with *α*_micro_ → ∞, to increase the probability of microgametes fusing with macrogametes and avert the low survival probabilities of microgametes under parthenogenesis. As this represents a situation in which macrogametes still fuse with microgametes (at a rate *α*_micro_/2) but do not fuse themselves, this can be biologically interpreted as the evolution of oogamy, with obligate sexual reproduction in the limit *α*_micro_ → ∞.

The exact stage along this branching that the population can reach depends on the maximum possible fusion rate, *α*_max_ and the cost to cell fusion, *C*. When amax is small (such that the proportion of fused daughter cells is small) isogamy can be stabilised. Meanwhile when the cost to cell fusion is very low (such that 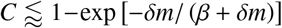, see Appendix C.4) the selective pressure for the macrogametes to reduce their fusion rate disappears (see Figure D.12), and the population evolves towards a state of pseudooogamy, in which both macrogametes and microgametes attract one another but with micogametes doing so at a faster rate [9].

### 3.3. In a switching environment without phenotypic plasticity, the population can evolve a bet-hedging strategy with isogamy at intermediate fusion rates

In Figure 4, we see that the analysis for the evolutionary dynamics in the case of a switching environment without phenotypic plasticity, Eq. (8), provides a good approximation to the dynamics of the full model (which accounts for multiple traits coexisting under a mutation-selection balance) realised via numerical simulation. We now see three broad evolutionary outcomes.

We begin by considering the intuitive limit of *τ*_1_ ≫ *τ*_2_. In this scenario the population spends almost all of the time in environment 1, and a comparatively insignificant amount of time in environment 2 (i.e. *P*_1_ ≈ 1 and *P*_2_ ≈ 0). Consequently, the population evolves approximately as if it were simply in a fixed environment with *β* = *β*_1_, and the conditions given for the fixed environment, Eq. (10), can be used to infer the evolutionary outcome. An analogous argument holds for the dynamics when *τ*_2_ ≫ *τ*_1_, but with *β* = *β*_1_ in Eq. (10). When the time spent in each environment is of the a comparable order however (e.g when *τ*_1_, and *τ*_2_ have not entirely dissimilar magnitudes), we find that the emergence of bet-hedging strategies.

Under a range of parameter conditions, we find analogous early evolutionary attractors for the fusion rate as in the fixed-environment case (see Eq. (10)), but with bet-hedging strategies for the daughter cell mass;

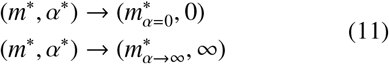

with

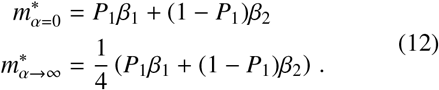

The population thus initially evolves either to high fusion rates or zero fusion rates, but with a mass which is the weighted average of the optimal strategy in either environment. Once on the 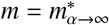 manifold towards the high-fusion rate attractor, evolutionary branching towards anisogamy and oogamy can occur as described in Section 3.2.

Under a more restricted set of parameter conditions however we find that a bet-hedging strategy for the fusion rate can also evolve; essentially a switching-induced fixed point can manifest, as illustrated in Figure 4. Here the tension between the evolutionary dynamics in the two environments (which can select for high fusion rate and large daughter cells in one environment, and zero fusion rate and small daughter cells in the other) can lead to the population being held in a state at which intermediate finite fusion rates and isogamy form the evolutionarily stable bet-hedging strategy.

In Appendix D.3, we show that the switching-induced fixed point is given by

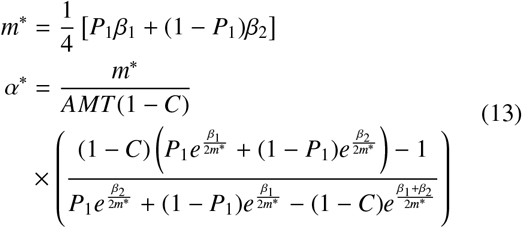

In Figure 4(c) we see that this switching induced fixed point is observed in the evolutionary simulations in which multiple traits can be held in the population under a mutation-selection balance. In the FRTI regime (gray lines), in which the environment changes more quickly, the population is held at the predicted mass *m*, while more variability between simulations is seen around the predicted fusion rate α as a result of the weaker selection on this trait. In the slower-switching FRTE regime (purple lines), a greater quantitative difference between the results of simulations and the analytic predictions is observed. We verify in Figure D.18 and Figure D.16 that this is not a result of evolutionary branching and that the population remains isogamous. However more importantly, the key prediction of finite fusion rate is indeed captured.

Biologically this bet-hedging strategy tells us that in a switching environment and when both sexual (cell-fusion) and asexual (non-cell-fusion) reproductive modes are possible, evolution can select for some investment in each to improve survival probabilities across the two environments.

### 3.4. In a switching environment with phenotypic plasticity, the population can evolve facultative sexual reproduction in response to harsh environments

Note that under the assumptions of costless and immediate phenotypic switching, the dynamics of (*m*_1_, *α*_1_) and (*m*_2_, *α*_2_) Eq. (9) are decoupled, significantly simplifying the analysis. The evolution of the traits in the respective environments are coupled however through the initial conditions from which they evolve, which must be the same (i.e. a phenotypically undifferentiated state). Although a large range of choices for these initial conditions could be argued, we here assume two parsimonious scenarios. In the first, we assume that the population has evolved to a stable non-fusing mass selected for in a single environment (e.g. 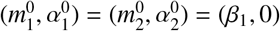); this is a scenario in which the alternate environment is in some sense novel and to which the population has not adapted. In the second scenario, we assume that the population has evolved to a bet-hedging strategy in mass (optimising the mass of daughter cells across the two environments), but has not yet evolved the capacity for binary cell fusion in either environment (e.g. 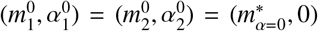, see Eq. (12)).

An illustrative phase portrait is shown in Figure 5. Consider the case in which a population has initially evolved under environment 1 to reach a stable state (*m*_1_, *α*_1_) = (*β*_1_, 0) (see red disk and surrounding purple circle, panel (a)). The population is now exposed to a second, harsher environment (*β*_2_ > *β*_1_) and allowed to evolve a phenotypically plastic response to this environment. Starting from an initial state 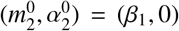) (see purple circle, panel (b)), traits *m*_2_ and *α*_2_ can evolve to (*m*_2_, *α*_2_) = (*β*_2_/4, ∞).

**Figure 5:**
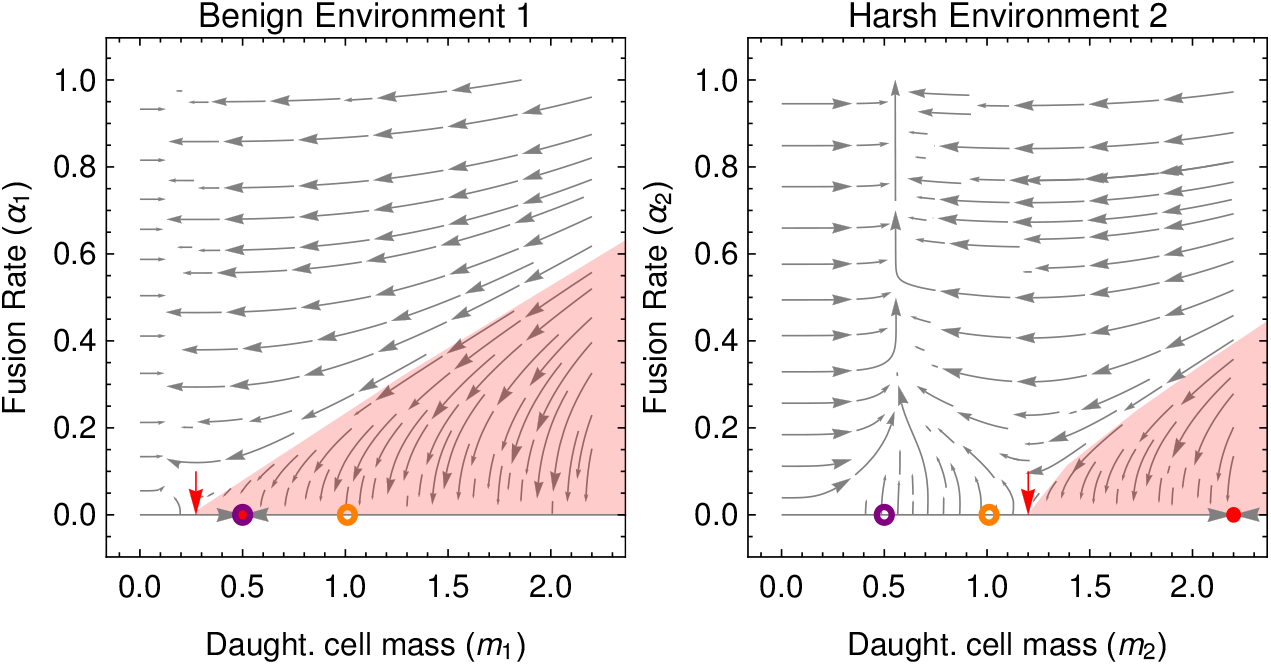
Illustrative phase portrait for co-evolutionary dynamics of (*m*_1_, *α*_1_, *m*_2_, *α*_2_) in a switching environment with phenotypic switching that exhibits facultative binary cell fusion. In both environment 1 (panel a) and environment 2 (panel b) the cost to cell fusion is *C* = 0.6, purple circles represent the initial condition 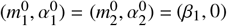, and orange cicles represent the initial condition 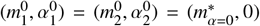, with 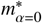 taken from Eq. (12). Environmental parameters are *β*_1_ = 0.5 and *β*_2_ = 2.2 making environment 1 the more benign envionment, in which the population typically spends a proportion *P*_1_ = 0.7 of its time. Remaining parameters are *A* = 100, *M* = 1 and *T* = 1.

Similarly, consider the case in which the population has initially evolved under the switching environments to reach a stable state bet-hedging strategy in mass 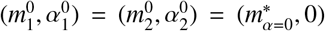 (see orange circles in Figure 5). Upon the evolution of phenotypic plasticity, in the benign environment we see the population traits relax to a stable state (*m*_1_, *α*_1_) = (*β*_1_, 0) (functionally asexual reproduction, see Figure 5(a)). However in the harsh environment, we see that the bet-hedging strategy in mass becomes unstable, and the population evolves towards traits (*m*_2_, *α*_2_) = (*β*_2_/4,0) in environment 2 (see Figure 5(b)).

In both scenarios described above, we see the emergence of facultative binary cell-fusion as a response to harsh environmental conditions that lower the survival probability of daughter cells. However we note that this is only possible if there is an appreciable decrease in survival probability (increase in resistance to survival, *β*) between the environments. In Figure 6, we summarise the key results over the *β*_1_ – *β*_2_ parameter plane. Here we assume that the cost to cell fusion is intermediate (1 – e^-2^ > *C* > 1 – e^-1/2^, i.e. 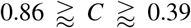) such that there are regions on the boundary *α* = 0 at which increased fusion rates are both selected for and against depending on the value of *m* (see Section 3.1 and Appendix E.2, Eq. (C.5)); this restricts us to the more interesting parameter regime in which different outcomes are possible in each environment. We further assume that the initial conditions are either at 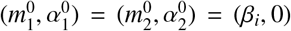 (where environment *i* is benign, such that *β_j_* > *β_i_*), or at 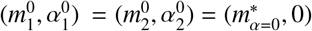 (where we note that 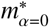 is a function of the probability of being in the benign environment, see Eq. (12)). We see that when one of the environments is not appreciably worse than the other, binary cell fusion does not involve in either environment. However when the difference between the environments grows more substantial, it is possible to evolve cell fusion in the harsher environment from initial condition 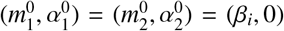 (where the population has first evolved towards the evolutionary optimum of the more benign environment). Finally when the difference between the environments is extreme, it is also possible to evolve cell fusion in the harsher environment from initial condition 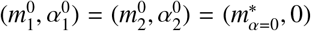 (where the population has first evolved towards a bet-hedging strategy in cell mass).

**Figure 6:**
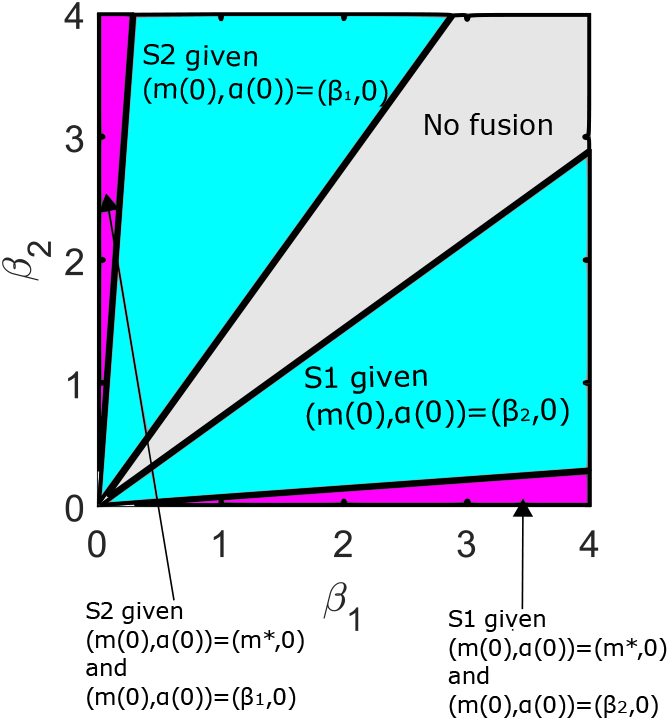
Regions in the *β*_1_-*β*_2_ parameter space where we observe sex in each of the two environments, given various initial conditions. S2 denotes the state of sex in environment 2 and S1 the state of sex in environment 1. Here, *C* = 0.5. The region plot is independent of *A*, *M* and *T*. *m*\* is the fixed point of a no-fusion population that undergoes bet-hedging in mass, defined by *m*\* = *P*_1_*β*_1_ + (1 – *P*_1_)*β*_2_ where *P_B_* = 0.7.

## 4. Conclusions

The evolution of sexual reproduction and its consequences for the subsequent evolutionary trajectory of populations is of general importance to biologists [18, 24, 30]. In this paper we have illustrated a reversal of the classic two fold cost of sex in organisms with distinct sexes [61]; in unicellular organisms without self-incompatibility systems (mating types) but with the capability for parthenogenic development, binary cell fusion is selected for even in the presence of substantial costs due to a two-fold survival benefit that comes from increased mass. The precise lessons theoretical lessons to take away from this study vary slightly depending on the how the model is interpreted.

Viewing our model as one for the evolution of eukaryotic sex in organisms capable of parthenogenesis, fusion rate can be straightforwardly be interpreted as fertilization rate. In a fixed environment, we find two broad evolutionary outcomes; selection for high fertilization rates (tending to obligate sexual reproduction) or selection for zero fertilization rates (functionally obligate asexual reproduction). To which of these attractors the population tends depends on the initial conditions and on the cost of cell-fusion, which in this context can be crudely interpreted as short term costs to syngamy and genetic recombination [24]. It is particularly interesting that the benefits conferred to cell-fusion through increased mass are sufficient to withstand remarkably high costs, with obligate sexuality remaining the only evolutionary attractor with costs equivalent to a loss of 39% of zygotes. In such a regime of extremely high fertilization success, our results indicate that initial evolutionary branching to anisogamy is a natural consequence, consistent with previous theoretical results [66]. However our results further show a route by which oogamy can also evolve from anisogamy in systems with parthenogenesis. Here sexual conflict emerges [67], whereby large daughter cells (e.g. macrogametes, which are capable of surviving without fusion) resist fusion with microgametes, but small daughter cells (e.g. microgametes, which are reliant on fusion with macrogametes for survival) evolve increasingly high fusion rates. We have seen that evolutionary progress along this isogamy-oogamy trajectory can be arrested at isogamy (if the maximum possible fusion rate is limited to a low value by environmental or energetic constraints) and at pseudooogamy (if costs to cell fusion are very low). It is also possible that additional modelling considerations known to stabilise these states, such as accounting for daughter cell-death during the fusion period [7], might have a similar effect here. Overall, the observed evolutionary states capture the full range of behaviours observed in species with parthenogenic gametes [15].

Perhaps most interesting is the case of switching environments with phenotypic plasticity. Here we find under a broad set of biologically reasonable conditions (costs to sexual reproduction equivalent to 39% - 86% zygote mortality and at least moderate changes in environmental quality) that high fusion rates are selected for in harsh environments and zero fusion rates are maintained in benign environments. This behaviour parsimoniously recapitulates the empirically observed reproductive strategies of numerous faculatatively sexual species, including *C. reinhardtii* [38], *S. pombe* [39] and *D. discoideum* [40]. This mechanism, under which cell fusion evolves to increase the survival probability of daughter (sex) cells, provides a complementary perspective on the frequent evolution of survival structures (resistant to environmental stress) that form following the formation of a zygote. These include ascospores in fungi [68] and zygote-specific stress-resistant stress wall in *C. reinhardtii* [69]. Note that such correlations between sexual reproduction and the formation of survival structures are not as easily explained under genetic explanations for the evolution of sexual reproduction, where engaging in both behaviours at once constitutes a simultaneous (and therefore potentially costly) change in genotype and temporal dislocation in environment [70, 71].

The results above are particularly interesting in the case of the evolution of early binary cell-fusion as a first step in the evolution of sexual reproduction. While most studies focus on the genetic benefits of cell-fusion [20] (including a functionally-diploid dikaryotic cell [26]), or the genetic benefits of mixed cytoplasm [30, 31] (which can also come with costs), the mechanism at play here is purely physiological. Yet, as addressed above, it naturally captures the empirical observation of binary cell-fusion as response to challenging environmental conditions. While the mechanism does not explain the evolution of sexual reproduction and genetic recombination itself, it does provide a nascent advantage to binary cell fusion that sets the stage for the evolution of sex by bringing nuclei into contact for prolonged periods, as well as countering short-term costs associated with recombination. Conceivably, if genetic recombination is beneficial for myriad genetic reasons in the long-term [23], it would seem natural that it would be instigated when the opportunity arises (i.e. when physiological survival mechanisms bring nuclei into close contact). In this scenario recombination may not be only a direct response to environmental variability [41, 43], but also to the correlated formation of a survival structure.

One element absent from our model is the fusion of multiple cells, which are likely to be selected for under the assumptions implicit in our model. There would clearly be an upper-limit on the number of fusions selected for, arising from the multiplicative effect of the fusion cost *C*, which would be present on each fusion. However in this context, it is interesting to note that one of the hypotheses for the evolution of self-incompatible mating types is as a signal to prevent the formation of polyploid cells [72]. Such a mechanism could also prevent the formation of trikaryotic cells should the cost of multiple fusions be too great. Thus, the model neatly preempts the second stage in models for the evolution of eukaryotic sex, the regulation of cell–cell fusion [25]. More generally, it is interesting to note that the conditions for facultative sexuality (e.g. harsh environmental conditions) broadly coincide with those for facultative multicellularity in both bacteria and eukaryotes, with starvation triggering the formation of fruiting bodies in myxobacteria [73, 74] and flocking in yeast [75, 76]. Meanwhile in *C. reinhardtii*, the formation of multicellular palmelloids and aggregates are an alternate stress response to sexual reproduction [77], as are the formation of fruiting bodies in *D. discoideum* [78]. In this multicellular context, the sexual behaviour of *D. discoideum* is particularly interesting, as once formed, the zygote attracts hundreds of neighboring cells that are then cannibalised for the provision of a macrocyst [79]. These various survival strategies are unified in our model as a mechanism for the evolution of binary cell fusion.

As addressed above, trade-offs on the evolution of cell-fusion rate, inbreeding, and the possibility of higher-order cell-fusions offer interesting avenues to extend this analysis. In addition, we have not accounted for the discrete nature of divisions leading to daughter cells, costs to phenotypic switching, non-local trait mutations, or pre-existing mating types. However perhaps the most obvious gap in our analysis is a mathematical characterisation of the branching from isogamy to anisogamy and oogamy. More generally, extending our mathematical approach leveraging adaptive dynamics to switching environments in other faculatatively sexual populations might prove particularly fruitful [80, 81].

In this paper we have extended the classic PBS model [3] in two key ways; allowing the fusion rate to evolve and subjecting the population to switching environments. In doing so, we have shown its capacity to parsimoniously capture the the evolution of obligate sexual reproduction, obligate asexual reproduction, faculative sexual reproduction, and environmentally triggered sexual reproduction in unicellular organisms. These results offer particularly interesting implications for the evolution of binary cell-fusion as a precursor to sexual reproduction, as well as suggesting common mechanistic links to the evolution of binary cell fusion. Moreover, our analysis emphasises the importance of exploring the coevolutionary dynamics of a range of evolutionary parameters and of developing mathematical approaches to deal with faculative sexual reproduction.

## Appendix A. Within generation dynamics

At the start of each generation, we assume a total of *A* unicellular organisms of mature cell size *M* divide to form daughter cells (or gametes). We further assume that the frequency of mutant adults in the population is given by 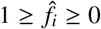, with *i* = *m* for mutants that change the mass of daughter cells and *i* = *α* for mutants that change the fusion rate of daughter cells.

### Appendix A.1. Fusion kinetics: mutant with different mass

If a mutation occurs changing the size of daughter cells produced by the mutant (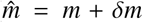 for resident daughter cell mass *m*), the fusion dynamics themselves (see Eq. (1)) are unaffected by the change in mass. However the number of daughter cells produced by the mutant will change. Denoting by *N* the number of resident daughter cells and 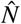 the number of mutant daughter cells, we have

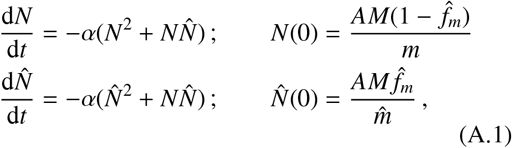

which has a solution

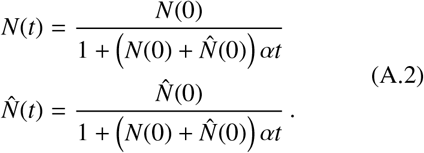

This allows us to determine the number of unfused cells of each type at the end of the fusion window at *U* = *N*(*T*) and 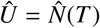, where we recall that *T* is the length of the fusion window.

We also need to determine the number of fused cells formed from the fusion of two residents, 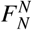, two mutants, 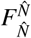, and a mutant and a resident, 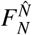. For this we need to solve

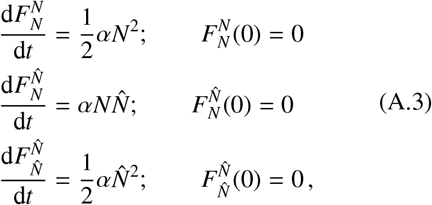

which can be solved by substituting for *N* and 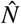 from Eq. (A.2) to yield

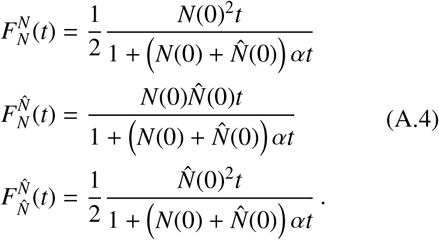

**Figure A.7:**
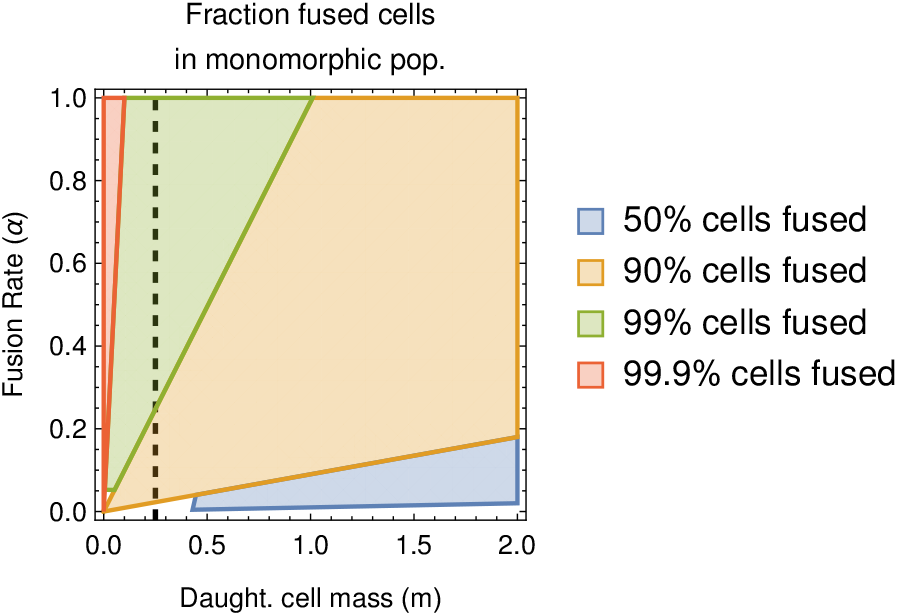
Illustration of the total proportion of cells that are fused at the end of a fusion period (length *T* = 1) in a monomorphic population (without branching) as a function of trait variables *m* and α. Parameters used are the same as those in 2. The vertical black dashed line gives the location of the manifold (*β*/4, *α*, along which the population is attracted when approaching the high-fusion attractor.

### Appendix A.2. Fusion kinetics: mutant with different fusion rate

If a mutation occurs changing the fusion rate of daughter cells produced by the mutant (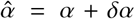 for residents with fusion rate α), the fusion dynamics themselves are altered relative to Eq. (1). We assume for simplicity that the fusion rate between resident-pairs is the mean of the fusion rate between the two types in isolation, such that

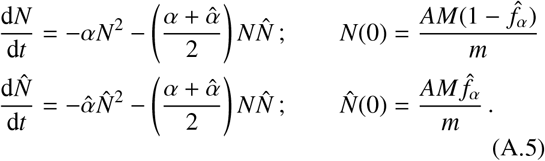

Solving this equation is slightly less straightforward than solving Eq. (A.1). However we can make analytic progress by making a change of variables and applying an approximation based on small mutational step size *δα*.

We introduce the transformed variables *N*_tot_ and 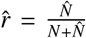, representing the total number of unfused cells and the frequency of unfused mutant cells respectively. Eq. (A.5) then becomes

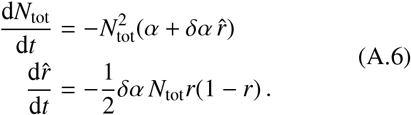

We now see that although this equation is also intractable, the leading order dynamics of *N*_tot_ are governed by *α*. Therefore when *α* ≫ *δα*, we make the approximation 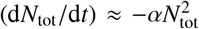 (we will return to the validity of this approximation as *α* → 0 later). We then

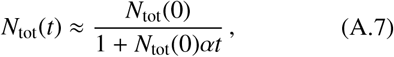

and substituting this into our equation for 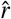 in Eq. (A.6), we can solve to obtain

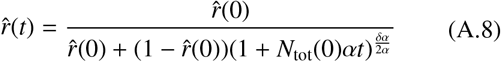

Inverting the transformation, we then arrive at

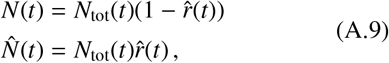

with *N*_tot_ and 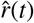 taken from Eqs. (A.7-A.8). We shall see in Appendix A.3 that calculating 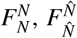, and 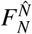 explicitly is in fact unnecessary, and these expressions for *N*(*t*) and 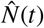 are sufficient for analytical progress.

### Appendix A.3. Change in mutant frequency over a generation: mutant with different mass

We begin by calculating the fitnesses of the resident and a mutant that changes the mass of daughter cells, *w_m_* and 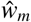 respectively, which are simply give by the total number of cells of each type at the end of a generation. Recalling that fused and unfused cells both survive with a probability governed by the parameter *β* and the mass of the cell *m_c_* (see Eq. (2)), and that fused cells survive with an additional probability (1 – *C*), we have

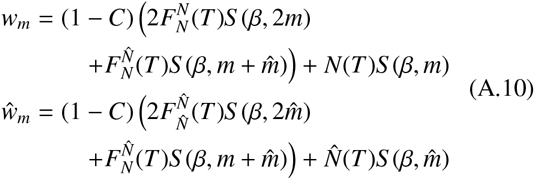

where 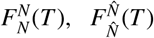, and 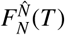 are taken from Eq. (A.4), and *N*(*t*) and 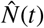 are taken from Eq. (A.2). These expressions can be used to calculate the frequency at the end of the generation, 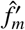, of a mutant that changes the mass of daughter cells as

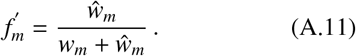

The change is the frequency of the mutant over the course of a generation is then 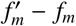.

### Appendix A.4. Change in mutant frequency over a generation: mutant with different fusion rate

Taking an analogous approach to Appendix A.3, we begin by calculating the fitnesses of the resident and a mutant that changes the fusion rate, *w_α_* and 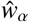 respectively. We obtain

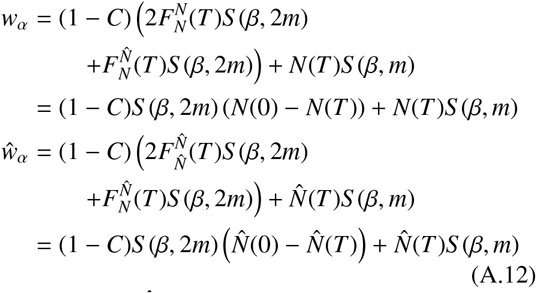

where *N*(*t*) and 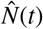 are now taken from Eq. (A.9). Here we have used the fact that since the survival function for fused cells, *S*(*β*, 2*m*), is independent of the composition of the fused cells (mutations here only affect fusion rate) the number of cell-types contributing to the fused cells can be inferred under cell conservation during the fusion period (e.g. for resident cell types 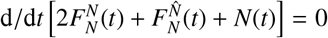, and similarly for mutant cell types).

These expressions can be used to calculate the frequency at the end of the generation, 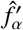, of a mutant that changes the fusion rate as

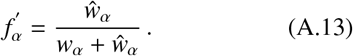

The change is the frequency of the mutant over the course of a generation is then 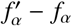.

## Appendix B. Invasion dynamics

Our aim is to derive the dynamics of the frequency of a mutant with a small mutation in either of the traits, *m* or *α*, over multiple generations. We introduce *t_g_* as a measure of the number of discrete generations.

**Figure B.8:**
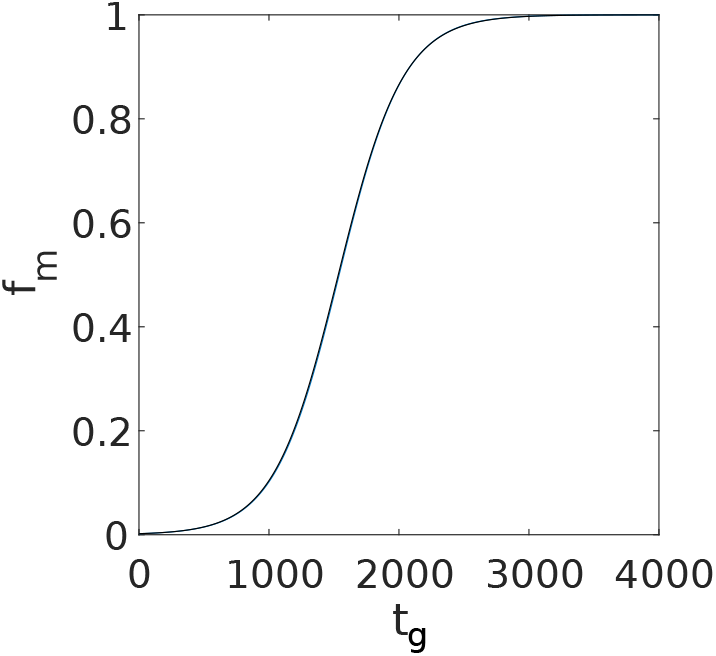
Invasion dynamics for a mutant with mass *m* + *δm*. Blue - analytical prediction using Eq. (B.1), black - numerical simulation. The initial condition is (*m*(0), *α*(0)) = (0.4, 0.1). Parameters are *δm* = −0.005, *f*_0_ = 0.002, *G* = 4000, *A* = 100, *M* = 1, *T* = 1, *C* = 0.6 and *β* = 1.

**Figure B.9:**
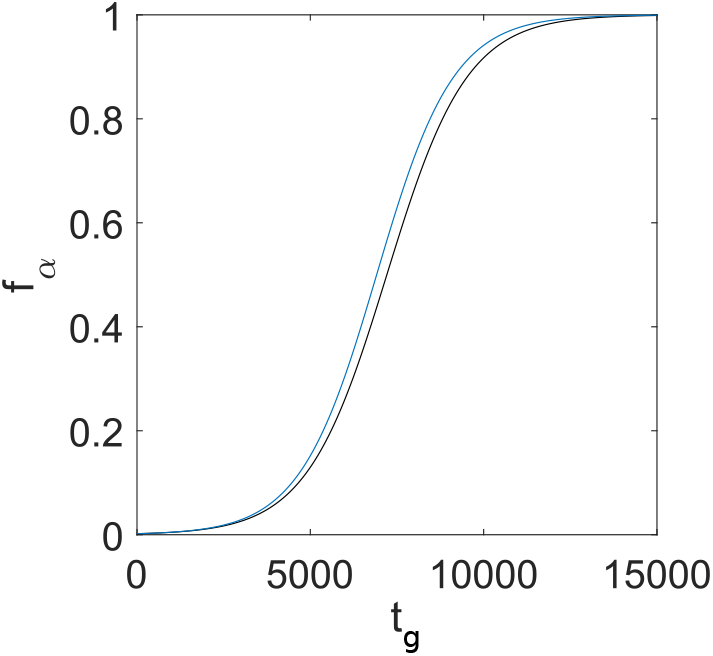
Invasion dynamics for a mutant with fusion rate *α* + *δα*. Blue - analytical prediction using Eq. (B.1), black - numerical simulation. The initial conditions and parameters are the same as in Figure B.2.

### Appendix B.1. Invasion ODE: mutant with different mass

We begin by deriving the dynamics for the frequency of a mutant that changes the mass of daughter cells. We begin by assuming that the mutational step size, *δm*, is small. Under these conditions, the frequency of mutants changes only by a small amount over the course of one generation, and we can approximate the frequency of mutants at the end of the generation (see Appendix A.3) by 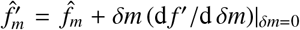, where 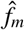 is the frequency of the mutants at the beginning of the generation. The dynamics of the mutant frequency over an invasion can then be approximated by

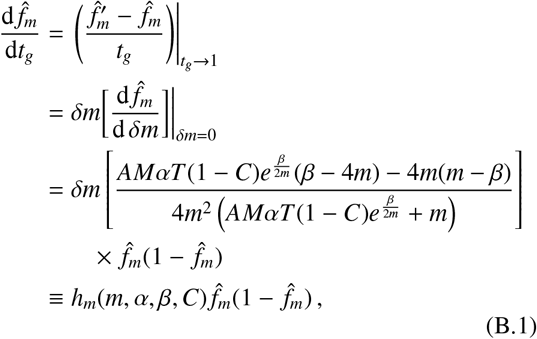

(see also Eq. (3)). We show in Figure B.8 that as expected, this is a good approximation for the dynamics when *δm* is small.

### Appendix B.2. Invasion ODE: mutant with different fusion rate

We now derive the dynamics for the frequency of a mutant that changes the fusion rate of daughter cells. Taking an analogous approach to Appendix B.1, we assume the mutational step size for mutation, *δα*, is small. We can then approximate the frequency of mutants at the end of the generation (see Appendix A.4) by 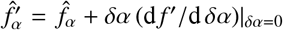, where 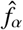 is the frequency of the mutants at the beginning of the generation. The dynamics of the mutant frequency over an invasion can then be approximated by

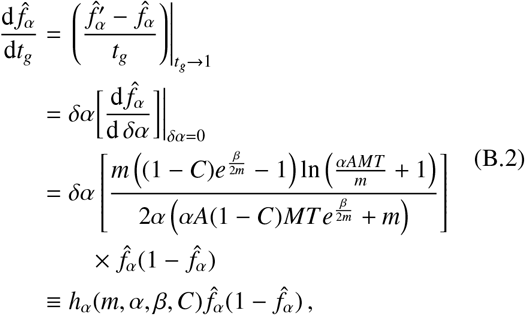

(see also Eq. (4)). We show in Figure B.9 that as expected, this is a good approximation for the dynamics when *δα* is small.

## Appendix C. Evolutionary dynamics: Fixed environment

### Appendix C.1. Derivation of evolutionary ODEs

We begin by noting that the functional form of 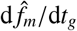 (see Eq. (B.1)) and 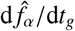 (see Eq. (B.2)) is 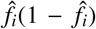, which implies a situation of trait substitution []; mutants are either driven to fixation or extinction, and polymorphic equilibria are not possible. This simplifies the subsequent analysis significantly.

Taking a classic adaptive dynamics approach [82] and define the invasion fitness of the mutants as their percapita rate of reproduction upon arising in the population (i.e. when 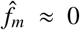 and 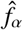). Under the standard assumptions of adaptive dynamics (i.e. that mutations are of small effect, 1 ≫ *δm*, *δα*, and occur sufficiently rarely that each mutation can fixate before a new mutation occurs), the evolutionary dynamics are given by

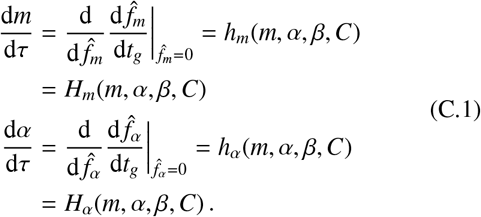

Substituting for 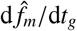 from see Eq. (B.1)), and 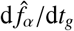 from Eq. (B.2), we obtain Eq. (5) in the main text.

### Appendix C.2. Analysis of ODEs

In this section we aim to analytically characterise the long term evolutionary behaviour of the population in a fixed environment, with dynamics given by Eq. (5), as illustrated in Figure 2.

We begin by calculating the evolutionary behaviour of *m* when the fusion rate is fixed to zero (*α* = 0). Solving *H_m_*(*m*, 0, *β, C*) = 0, for 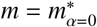 we obtain

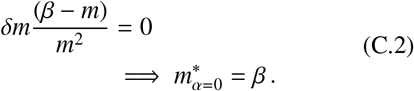

This fixed point is always stable in the *m*-direction when *α* = 0. When modelling the evolution of early cell fusion, the state 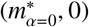 arguably serves as a reasonable initial condition for the coevolution of *α* and *m*, under the assumption that the evolutionary dynamics of daughter cell mass preceded the evolution of the physiological machinery required binary cell-fusion.

We now turn to the evolutionary behaviour of *m* and *α*. For the zero-fusion point 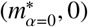 identified above to remain stable once evolution on α is allowed requires that [d*α*/d*τ*] |_(*m*=*β,α*=0)_ < 0 (i.e. that evolution selects against increases in *α*). Stating this condition in full, we have

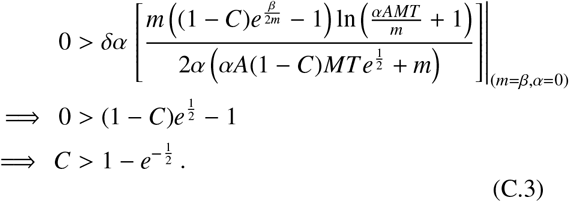

If fusion costs exceed this this value of *C* ≈ 0.39, evolutionary trajectories starting at 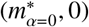 will remain there (see Figure 2, panels b-c). Conversely if fusion costs do not exceed *C* ≈ 0.39, evolutionary trajectories starting at 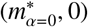 will initially experience a selective pressure for increasing *α*, due to the fact [d*α*/d*τ*]|_*m*=*β*_ > 0 under these conditions (see Figure 2, panel a); the dynamics are then pulled to another attractor, which we characterise below.

When *C* > 1 – exp (–1/2) (but less than another critical value, yet to be determined), only a subset of initial conditions fall within the basin of attraction of this attractor described above (see Figure 2, panel b), with remaining initial conditions leading to an evolutionary state at which *α*\* → ∞. Conversely, when *C* < 1 – exp (−1/2), *all* initial conditions lead to this attractor at which *α*\* → ∞ (see Figure 2, panel a). We now calculate the mass to which the population evolves at this second attractor. Taking the limit *α* → ∞ in the evolutionary dynamics for *m* (see Eq. (5)) and solving for zero;

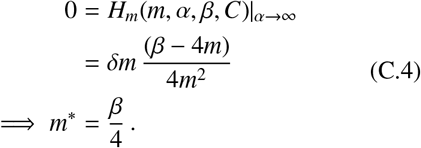

Therefore the second early evolutionary attractor is at (*m, α*) → (*β*/4, ∞).

For even greater costs of fusion, *C*, our mathematical analysis (which assumes monomorphic resident populations and no evolutionary branching) suggests that 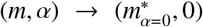 becomes the only attractor, with 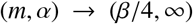 ceasing to be an attractor. To determine the critical cost at which this occurs, we take [d*α*/d*τ*]_*m*=*β*/4_ and calculate the conditions under which this is negative when *α* is large (i.e. when (*m, α*) → (*β*/4, ∞) is no longer attracting, but repelling). Expanding [d*α*/d*τ*]_*m*=*β*/4_ in small 1/*α*, we find that to leading order we must have;

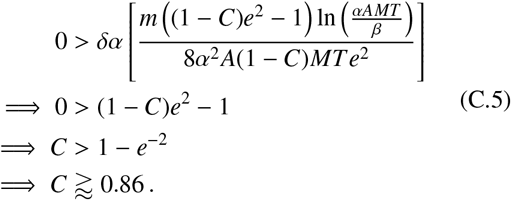

However, while this accurately captures the short-term evolutionary dynamics, we again see evolutionary branching (which our model does not account for) should trajectories approach the *m* = *β*/4 manifold. Unless costs are exceedingly high (*C* ≈ 1), this eventually leads to obligate oogamous sexual reproduction (with sexual conflict driving microgametes to increase their fusion rates to macrogametes at extremely high rates). We discuss this more in Appendix C.4.

### Appendix C.3. Implementation of simulations

Here we detail the process of numerical simulation. We employ a multistrain model whereby a mutation occurs at rate *μ*. This rate is the inverse of the expected number of fusion processes (i.e. generations) until the next mutation event. Before each successive mutation event, the number of fusion processes until the next mutation event is determined by generating a number from a *Geo*(*μ*) distribution and taking the inverse of that number. In an *S* strain model, if a given strain *i* has a trait value of (*m_i_*, *α_i_*) and has frequency *f_i_*, then the mean population trait value is given by

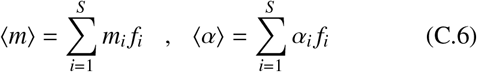

The simulations in Figure 2 are repeated for 2500 mutation events. Below, we detail how we simulate the dynamics on each timescale.

#### Fusion Kinetics

We construct our model for fusion kinetics assuming that each strain characterised by their unique (*m_i_, α_i_*) can fuse with one another. The input parameters of this function are the trait values of each strains ***m*** and ***α***, their frequency in the preceding adult generation ***f***, and the parameters *A, M* and *T*. The fusion kinetics simulations are then run according to

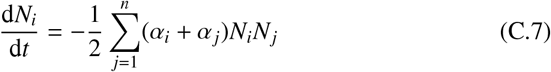

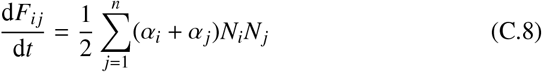

where *F_ij_* is the number of fused cells formed from the fusion of strain *i* and *j*, and initial conditions

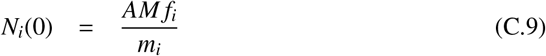

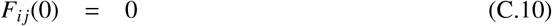

The fusion process is run for a fixed time period *T*. At the end of this time period, the function outputs the number of cells of each type. These include the number of unfused cells with each trait pair, *N_i_*(*T*), and fused cells of each type *F_ij_*(*T*).

#### Single Generation Dynamics

In this simulation, we simulate the frequency of each strain after the end of each generation taking into account the survival probability of each strain. Upon maturation, only a fraction of unfused and fused cells survive to adulthood. We use the outputs of the fusion kinetic function along with the Vance survival functions to calculate the probability that each progeny survives into adulthood. The input parameters of this function are *N_i_*(*T*) and *F_ij_*(*T*), ***m***, *α, β* and *C* whilst the output is ***f***, the frequency of all strains at the end of the generation.

#### Invasion Dynamics

The invasion dynamics are run for approximately one unit of *τ* (i.e. until the next randomly chosen mutation event after *G* generations). As outlined in the opening paragraph of Appendix C.3, *G* is generated from a *Geo*(*μ*) distribution following each mutation event. The inputs of this function are ***f***, ***m***, *α, A, M, T, C, G* and *β* and the output is the frequency of each strain ***f*** after one unit of *τ* (i.e after *G* fusion processes). Please note we have adopted one single notation for the frequency of each strain.

#### Evolutionary Dynamics

The evolutionary dynamics have the input parameters *δ, μ*, ***f***, *A*, *M*, *T* and *C* where *δ* is the mutational stepsize, the initial frequency of a newly introduced mutant *f*_0_ is chosen to be small (0.002 in our model). We initialise the simulation with two strains, where each strain is characterised by a unique pair of trait values e.g. (*m_i_, α_i_*) for strain i. A mutant is introduced into the population after a random number of fusion processes *G* (generated from a *Geo*(*μ*) distribution) at frequency *f*_0_. The mutant is chosen to occur in either α or *m* with equal probability 1/2. The mutation also acts to increase/decrease the trait value each with probability 1/2. Upon introduction of this mutant, the invasion dynamics of the population is run for *G* fusion processes. In the meantime, the mean mass and fusion rate of the population is recorded using Eq. (C.6). Next, we repeat the process of introducing a new mutant into the population. Since the population now has more than two strains, the strain that mutates is chosen with probability weighted by the frequency of each strain. There is now the possibility of back mutation to one of the existing strains. In this case, if *f_k_* is frequency of this strain before the introduction of the phenotypically identical mutant, then the frequency of this strain becomes (1 – *f*_0_)*f_k_* + *f*_0_ after its introduction. Care is taken when choosing the value of *μ*, since if a mutation occurs after a large number of fusion processes, fixation is likely to occur, reducing the model’s sensitivity to selection strength. 30 sample realisations of the stochastic model is obtained.

### Appendix C.4. Evolutionary Branching in Daughter Cell Mass and Fusion Rate

In this section we present additional numerical results investigating the evolutionary branching that occurs in simulations on the manifold *m* = *β*/4 (along which d*α*/d*t* ≈ 0).

#### *In a fixed environment, when α*_max_ *is small, branching in mass does not occur*

When the maximum fusion rate, *α*_max_, is very limited (e.g. by turbulence) we find evolutionary branching in daughter cell mass does not occur and the population is held in an isogamous state. This behaviour is illustrated in Figure D.14.

#### In a fixed environment, when C is small, branching in fusion rate still occurs but no longer gives rise to oogamy

When the cost to cell fusion, *C* is very small, we find that although branching in both daughter cell mass and fusion rate still occurs, the branching in α no longer acts to decrease the fusion rate of macrogametes. Therefore, we observe pseudooogamy but not oogamy. This behaviour is illustrated in Figure D.12. We can estimate the critical value of *C* at which this occurs by comparing when the survival probability of a macrogamete that does not fuse with a microgamete with that of a macrogamete that does fuse with a microgamete at a cost; if this first probability exceeds the second, there should be no evolutionary pressure for oogamy to evolve. We find

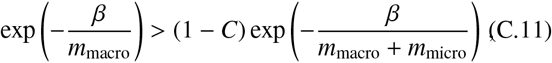

or, noting that *m*_macro_ approaches *β* and *m*_micro_ approaches *δm*

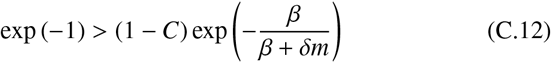

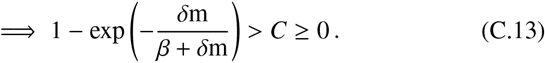

#### In a fixed environment, when C is very large, branching in mass and fusion rates can still occur, with obligate oogamous sexual reproduction possible (dependent on initial conditions)

Our analysis of the dynamics of Eq. (5) in Appendix C.2 suggested that zero fusion rates (asexuality) is the only evolutionary attractor when costs to binary cell fusion exceed *C* ≈ 0.86. However, simulations reveal that although d*α*/d*t* < 0 along the *m* = *β*/4 manifold, any trajectory that approaches this manifold can experience branching in daughter cell mass. Once branching in cell-mass occurs, the smaller daughter cells (e.g. microgametes) again experience a strong selective pressure to increase their fusion rate, despite the high costs imposed by cell-fusion; sexual conflict drives the population evolves towards obligate oogamous sexuality imposed by motile microgametes. Thus as for *C* < 1 there are always a subset of initial conditions that lead to trajectories along the *m* = *β*/4 manifold (most obviously initial conditions on the manifold itself) obligate sexual reproduction thus remains one of the two evolutionary outcomes, albeit requiring an increasingly small and biologically unrealistic set of initial conditions. Even up to significantly high costs, we see initial conditions that can allow the system to evolve to obligate sexuality e.g. *C* = 0.99 (see Figure D.19). When *C* = 1, this possibility disappears, and zero fusion rates (obligate asexuality) are uniformly selected for. This behaviour is illustrated in Figure D.20.

## Appendix D. Evolutionary dynamics: switching environments without phenotypic plasticity

We first tackle the derivation of the approximate dynamics in the FRTI (fast relative to invasion) switching regime in Appendix D.1, before verifying the qualitative robustness of these results in the FRTE switching regime in Appendix D.1.

### Appendix D.1. Derivation of evolutionary ODEs: FRTI

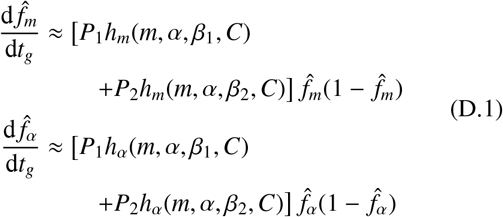

where *h_m_*(*m,α,β, C*) and *h_α_*(*m,α,β, C*) are given in Eq. (B.1) and Eq. (B.2) respectively. In Figure D.10 we show that this indeed is a good approximation of the dynamics when *δm* and *δα* are small and when the residency times in each environment are small relative to the invasion time. Note that as in the case of the fixed environment, the functional form of *f_i_* in these equations implies that trait substitution occurs for independent mutations on the daughter cell mass and fusion rate.

We now apply the same approach as in Appendix C.1 (see Eq. (C.1)) to obtain

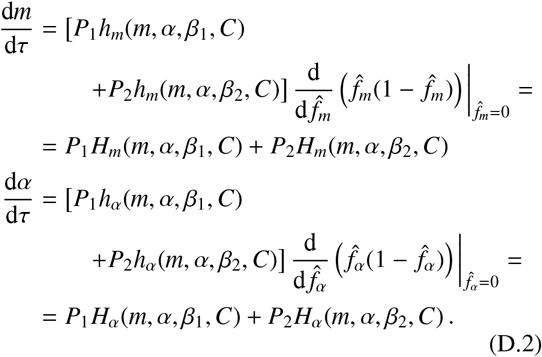

We see in Figure 4 that these also provide a good approximation of the evolutionary dynamics.

### Appendix D.2. Derivation of evolutionary ODEs: FRTE

In the FRTI scenario in the previous section, we supposed that switching between the environments was happening sufficiently regularly relative to the timescale of invasion that the effective invasion dynamics could be described by a weighted mean of the invasion dynamics in both environments (see Eq. (D.1)). Applying an analogous logic, we now assume in the FRTE scenario that switching between the environments occurs regularly relative to the timescale of evolution (the arrival rate of new mutations) that the effective evolutionary dynamics can be described by a weighted mean of the evolutionary dynamics in both environments; that is

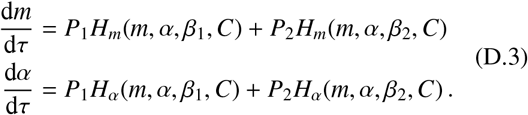

**Figure D.10:**
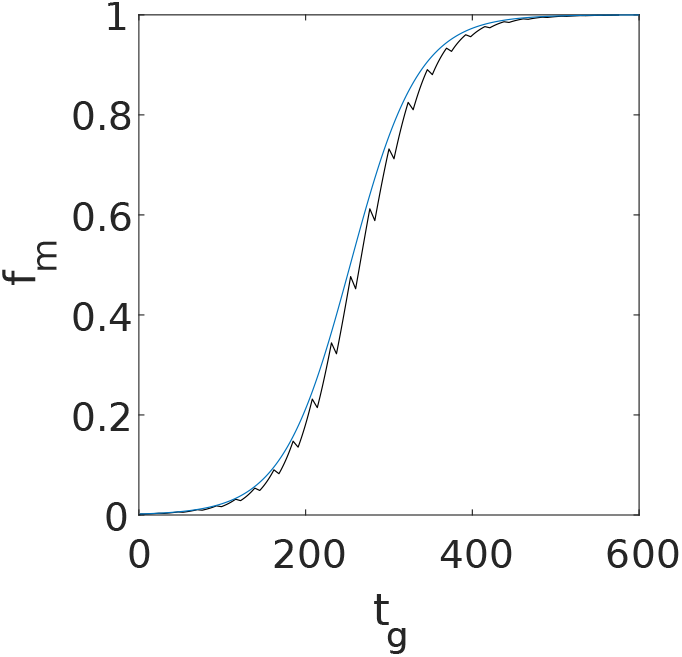
Invasion dynamics for a population undergoing bet-hedging when the environment switches FRTI. Blue curve is the analytical approximation using Eq. (D.1) and jagged curve is the numerical simulation. Mutation occured in mass with *δ* = 0.005, *f*_0_ = 0.002, (*m*(0), *α*(0)) = (0.3, 0.1), *λ_GB_* = 1/6, *λ_BG_* = 13/222. All other parameters the same as Figure 4.

We note that these are in fact exactly the same evolutionary dynamics as derived in the FRTI scenario (see Eq. (D.3)). In Figure 4 we show that these do indeed remain a good approximation for the dynamics in the FRTE scenario. The difference between the dynamics in both regimes is quantitative, rather than qualitative. In the very-fast switching FRTI regime, the populations follow the effective dynamics very closely. In the comparatively slower FRTE regime, although the populations no longer follow the dynamics as well, the qualitative picture of the dynamics is still captured by (D.3). In particular, we still observe an evolutionary stable state of intermediate fusion rate (see Figure 4).

### Appendix D.3. Analysis of ODEs

We begin, as in Appendix C.2, by considering the evolutionary behaviour of *m* when the fusion rate is fixed to zero (*α* = 0). We recall that if the population was fixed in environment 1 or 2 with *α* = 0, the mass of daughter cells would evolve to *m* = *β*_1_ and *m* = *β*_2_ respectively. In the switching environment we instead find the bet-hedging strategy

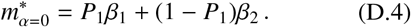

Meanwhile the region of the boundary *α* = 0 over which reduced fusion rates are selected for is given by [d*α*/d*τ*]|_*α*=0_ < 0, or

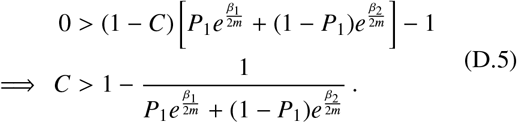

If this condition holds for 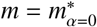, then the switching-induced fixed point 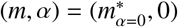 is stable.

We next turn to the high fusion rate attractor, for which *α* → ∞. In a similar manner to Eq. (C.4) (albeit with *H_m_*(*m,α,β, C*) replaced with *P*_1_ *H_m_*(*m, α, β*_1_, *C*) + *P*_2_*H_m_*(*m, α, β*_2_, *C*)), we find the bet-hedging strategy

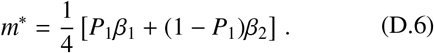

However, unlike in the fixed environment case, (*m, α*) = (*m*\*, ∞) is not always an evolutionary attractor.

In the switching environment, a third evolutionary attractor can emerge, brought about by a balance between selection for high fusion rates in one environment and zero fusion rates in the other. Solving *P*_1_ *H_α_*(*m, α, β*_1_, *C*) + *P*_2_*H_α_*(*m, α, β_2_, C*) = 0 for α we obtain

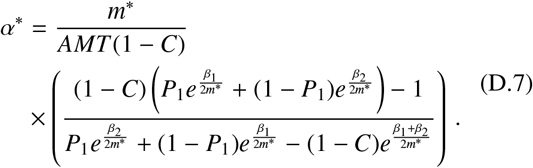

A good approximation for *m*\* in this equation can be deduced by noting that the fixed point sits on a vertical manifold of trajectories along which *m* is approximately held constant as *α* → ∞; thus we substitute ≈ *m*\* from Eq. (D.7) to obtain the approximate expression for the attractor (*m*\*,*α*\*) given in Eq. (13). In Appendix D.5, we verify from simulations that evolutionary branching does not occur at this switching-induced fixed point, and that the population is instead held in a state of isogamy.

### Appendix D.4. Implementation of simulations

In the bet-hedging scenario, we simulate the evolutionary dynamics by implementing a Gillespie algorithm. In particular, we introduce mutations and environmental switching events randomly with geometrically distributed waiting times, where the waiting time is measured in units of number of fusion processes *t_g_*. To simulate the FRTI regime, we set the environmental switching rates to larger values than the mutation rate i.e. *λ_GB_, λ_BG_* >> *μ*. Likewise, in the FRTE regime, we set *λ_GB_* and *λ_BG_* to smaller values than *μ*. The mutation rate in the numerical simulation of the FRTI regime overlaid in Figure 4 (a) is *μ* = 3.5 × 10^-4^ and the switching rates are *λ_GB_* = 67/532 and *λ_BG_* = 1/4. For Figure 4 (b) we have *μ* = 3.5 × 10^-4^ and the switching rates are *λ_GB_* = 1/6 and *λ_BG_* = 13/222. For the FRTE regime, *μ* = 5 × 10^-4^ and the switching rates are 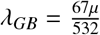 and *λ_BG_* = *μ*/4 in Figure 4 (a) and *λ_GB_* = *μ*/6 and 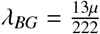 in Figure 4 (b).

### Appendix D.5. Absence of Evolutionary Branching at Switching-Induced fixed point

In this section we present additional numerical results that confirm that evolutionary branching does not occur at the switching-induced fixed point calculated in Eq. (13). We see that under both FRTI and FRTE, the population is held in a state of isogamy. This behaviour is illustrated in Figure D.15 to Figure D.18.

## Appendix E. Evolutionary dynamics: switching environments with phenotypic plasticity

### Appendix E.1. Derivation of evolutionary ODEs

With phenotypic plasticity, the evolutionary trajectory of the population in one environment is independent of that in the other. A mutation would exert a selection pressure in only one environment. If a mutation occurs in environment *i*, the invasion dynamics is given by

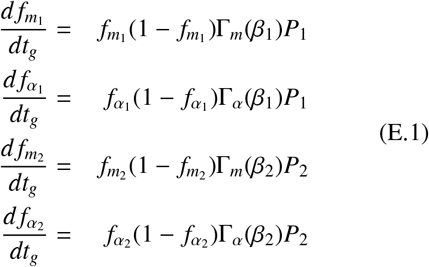

from which we can apply the same approach as Appendix C.1 to obtain the evolutionary dynamics given by Eq. (9).

### Appendix E.2. Analysis of ODEs

In the case where 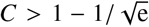, there exists a scenario whereby obligate syngamy would evolve if the initial state of the population is (*β*_1_, 0). The condition for obligate syngamy to evolve in environment 2 is for 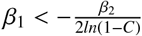. Due to the condition 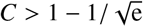, obligate syngamy does not evolve in environment 1. If the population starts at (*β*_2_, 0) on the other hand, obligate syngamy evolves in environment 1 if 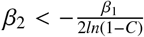. If the initial point of the population is (*m*\*, 0) where *m*\* = *P*_1_*β*_1_ + (1 – *P*_1_)*β*_2_, obligate syngamy evolves in environment 1 if 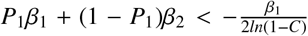 provided that *β*_1_ > *β*_2_. For the same initial point, obligate syngamy evolves in environment 2 if 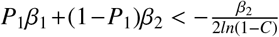 provided that *β*_2_ > *β*_1_.

**Figure D.11:**
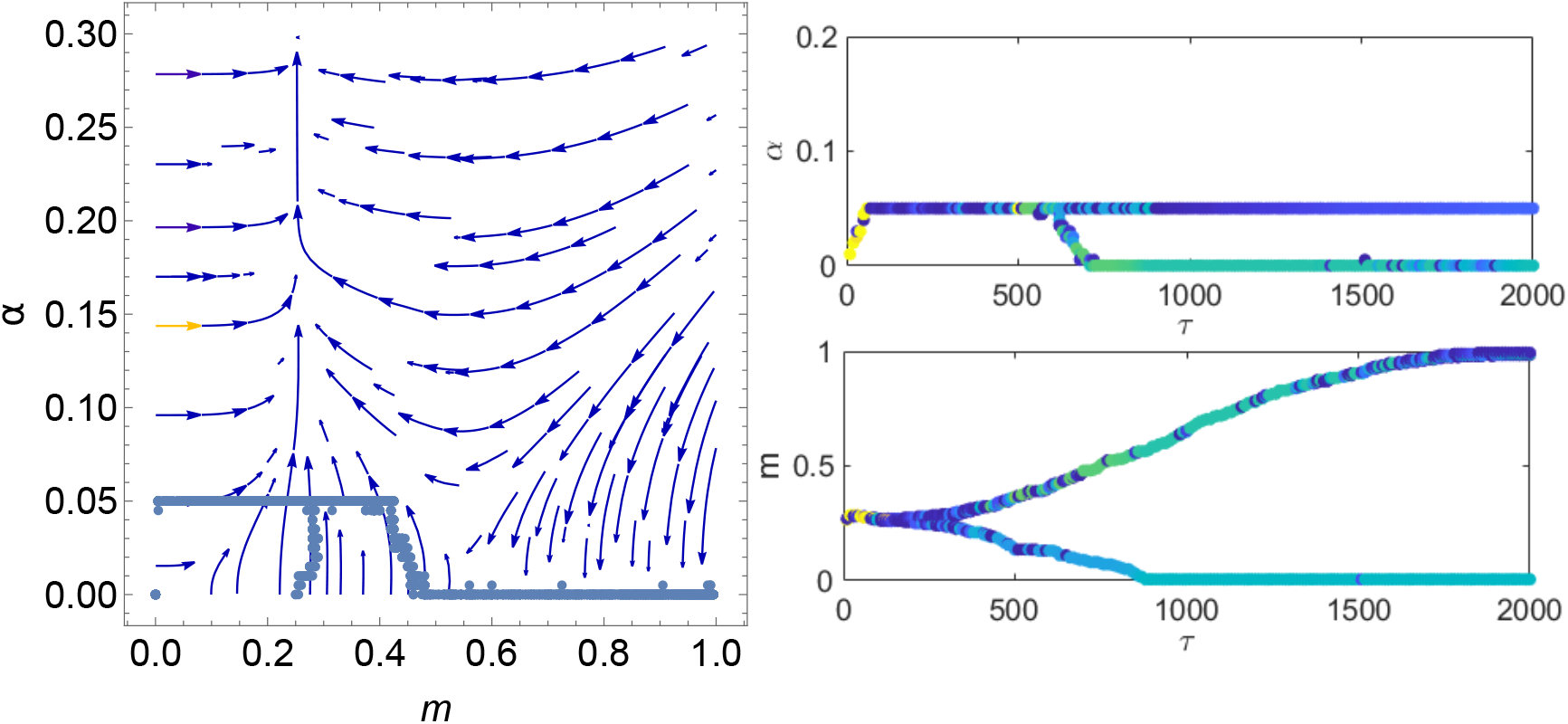
Numerical illustration of evolutionary branching of Figure 2 (b) with *α_max_* = 0.05. All parameters same as Figure 2 (b) except (*m*(0), *α*(0)) = (0.35, 0) and simulation run for 2000/*μ* generations.

**Figure D.12:**
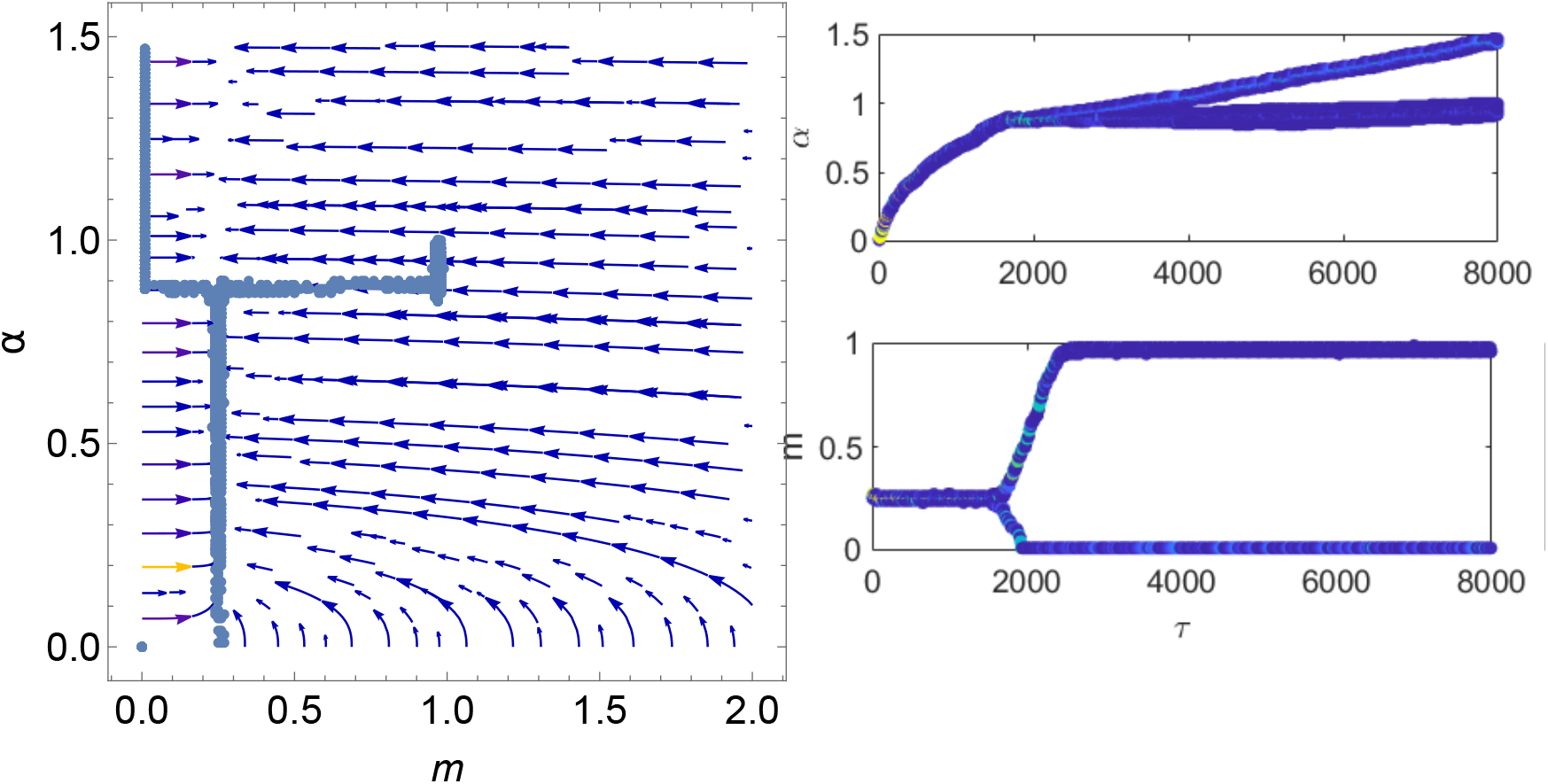
Numerical illustration of evolutionary branching for the case where *C* = 0. All other parameters the same as Figure 2, except (*m*(0), *α*(0)) = (0.25, 0) and simulation run for 8000/*μ* generations. Here, we observe no oogamy as the fusion rates of large daughter cells (macrogametes) do not tend to 0.

### Appendix E.3. Implementation of simulations

We numerically simulate the evolutionary dynamics starting from the initial states as outlined in Appendix E.1. To simulate the evolutionary dynamics for a population with initial conditions (*m*(0), *α*(0)) = (*β*_1_, 0), we use a fixed value of *β*_2_, and run the evolutionary dynamics as described in Appendix C.3, for various values of *β*_1_ (ranging from 0 to 2 in our simulations). To simulate the evolutionary dynamics for a population with initial condition (*m*(0), *α*(0)) = (*P*_1_*β*_1_ + (1 – *P*_1_)*β*_2_, 0), we again run the evolutionary dynamics as described in the above sentence, using the same fixed value of *β*_2_. The parameters used to run the evolutionary dynamics are *δ* = 0.02, *μ* = 1/500, *f*_0_ = 0.002 and simulation run for 400/*μ* generations. All the other parameters as in Figure 6. Running it for 400/*μ* generations provides sufficient time for the stochastic trajectory to reach the |*m* ≈ *β*/4| manifold in the obligate fusion environment. At the 400th mutation event, we check whether the conditions |*m*–*β*/4| < *δ* and *α* > *δ* are satisfied to determine whether the trajectory has reached the *m* ≈ *β*/4 manifold. A numerical simulation is shown in Figure E.21.

**Figure D.13:**
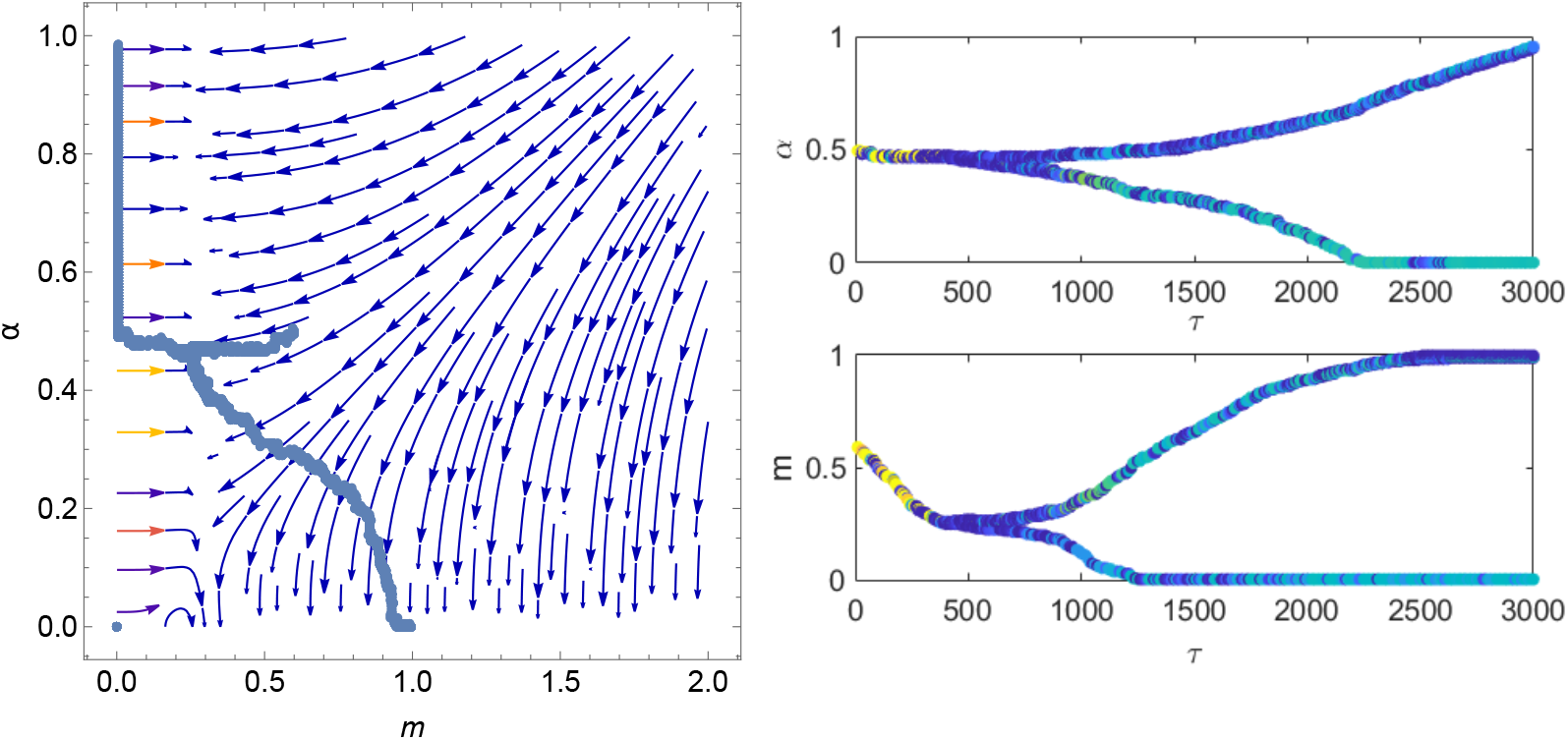
Numerical illustration of evolutionary branching for the case where *C* = 0.9. All other parameters the same as Figure 2, except (*m*(0), *α*(0)) = (0.6, 0.5) and run for 3000/*μ* generations.

**Figure D.14:**
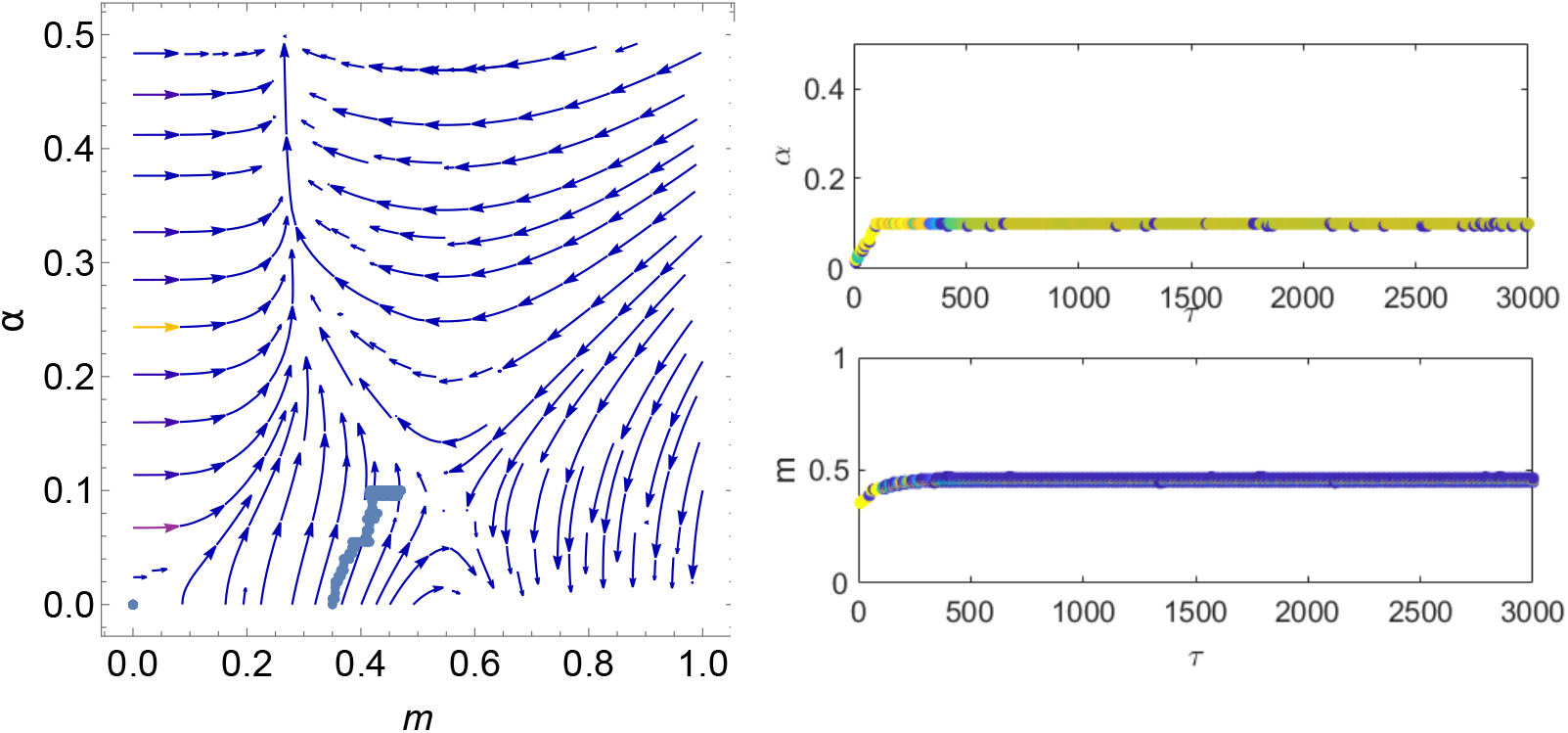
In the case where *α_max_* = 0.1 and *T* = 0.1 i.e *α_max_ T* = 0.01 with all other parameter conditions same as Figure 2 (b), we observe no evolutionary branching. Given that *C* = 0.6, *A* = 100, *M* = 1 (conditions of Figure 2), branching is absent for *α_max_, T* ≤ 0.01. This amounts to 69% of cells fused in each fusion generation.

---

**Figure D.15:**
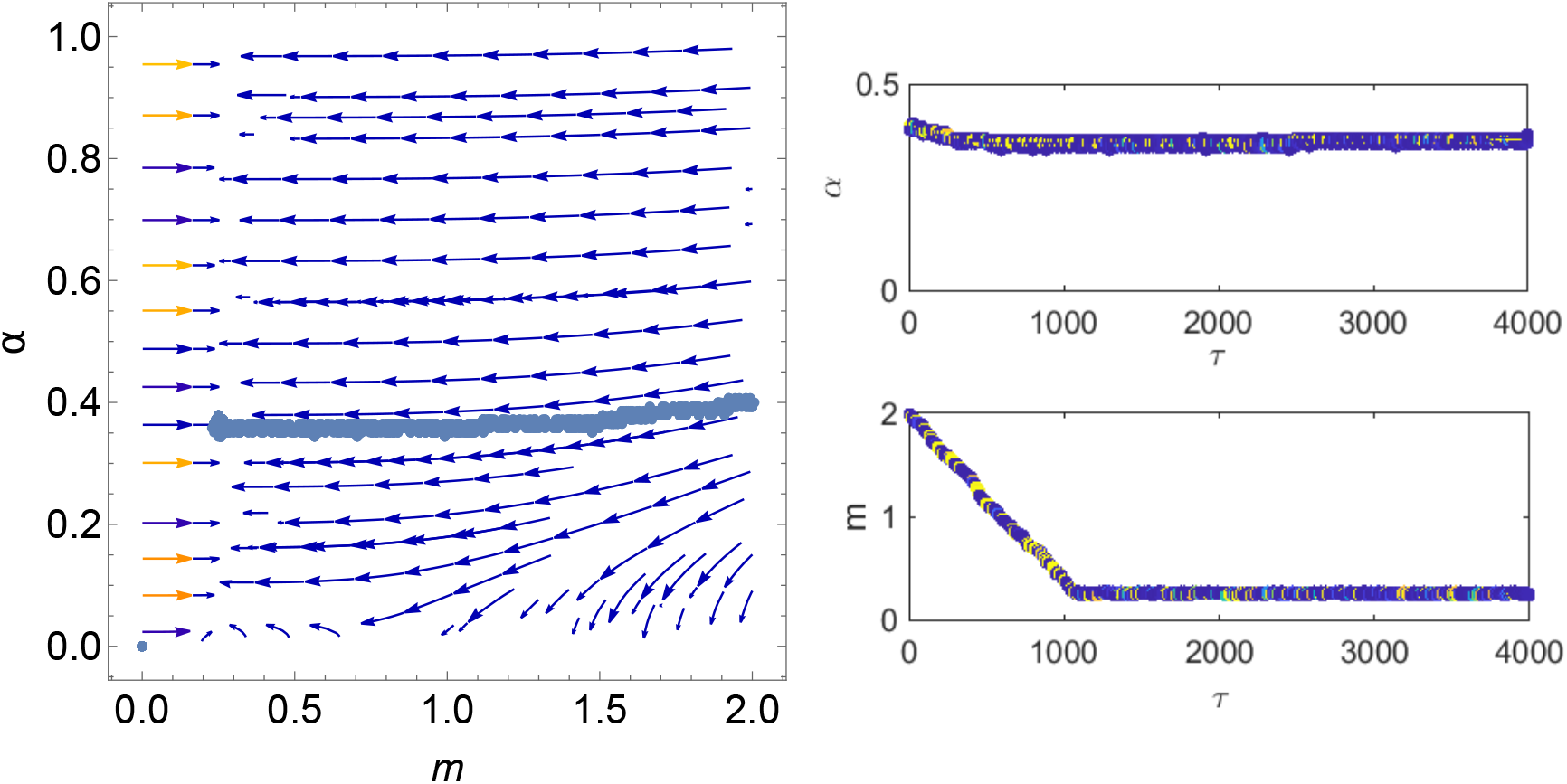
Numerical illustration showing an absence of evolutionary branching for a system undergoing bet-hedging in an environment that switches FRTI. All other parameters the same as Figure 4 (a) and the system is run for 4000/*μ* generations.

**Figure D.16:**
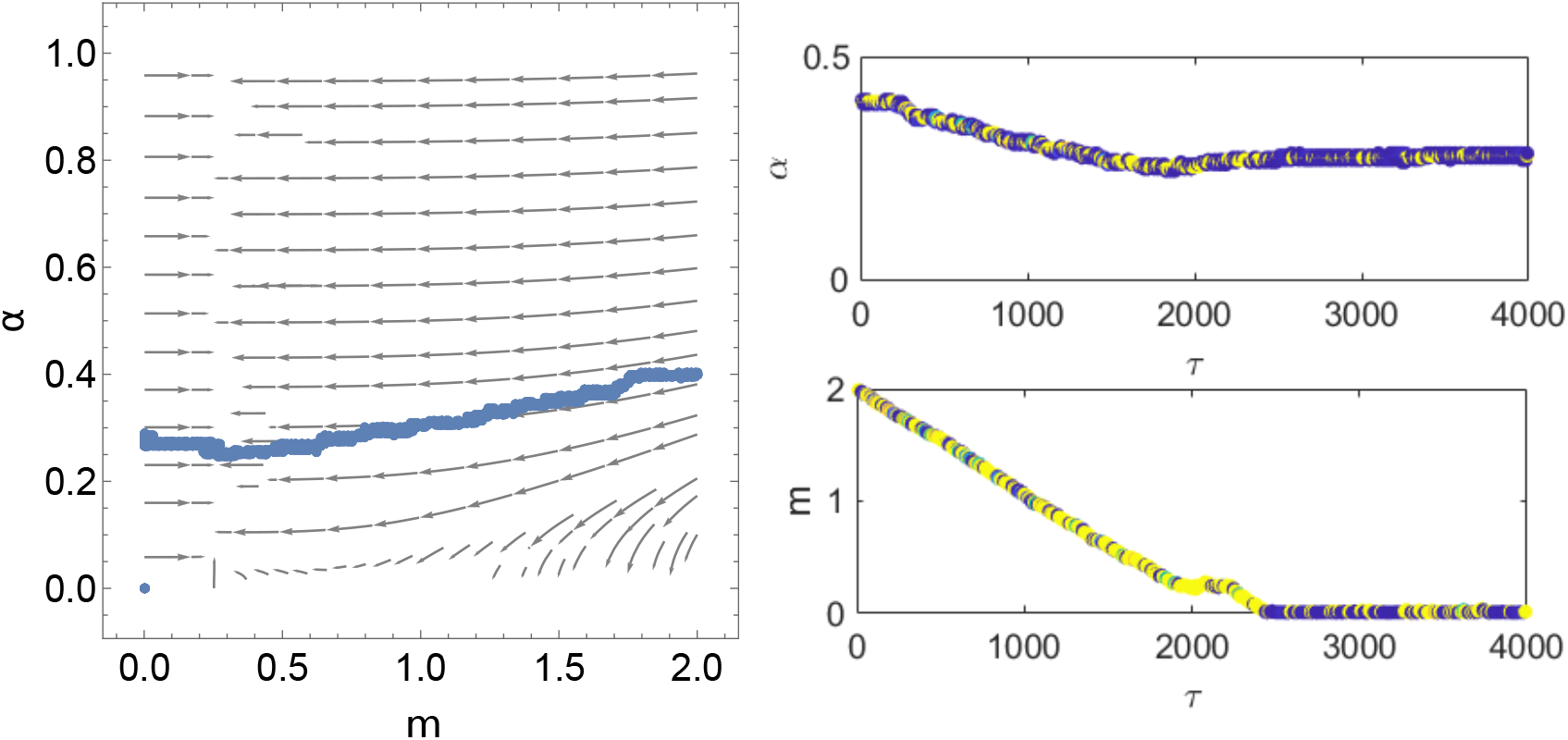
Numerical illustration showing an absence of evolutionary branching for a system undergoing bet-hedging in an environment that switches FRTE. All other parameters the same as Figure 4 (a) and the system is run for 4000/*μ* generations.

**Figure D.17:**
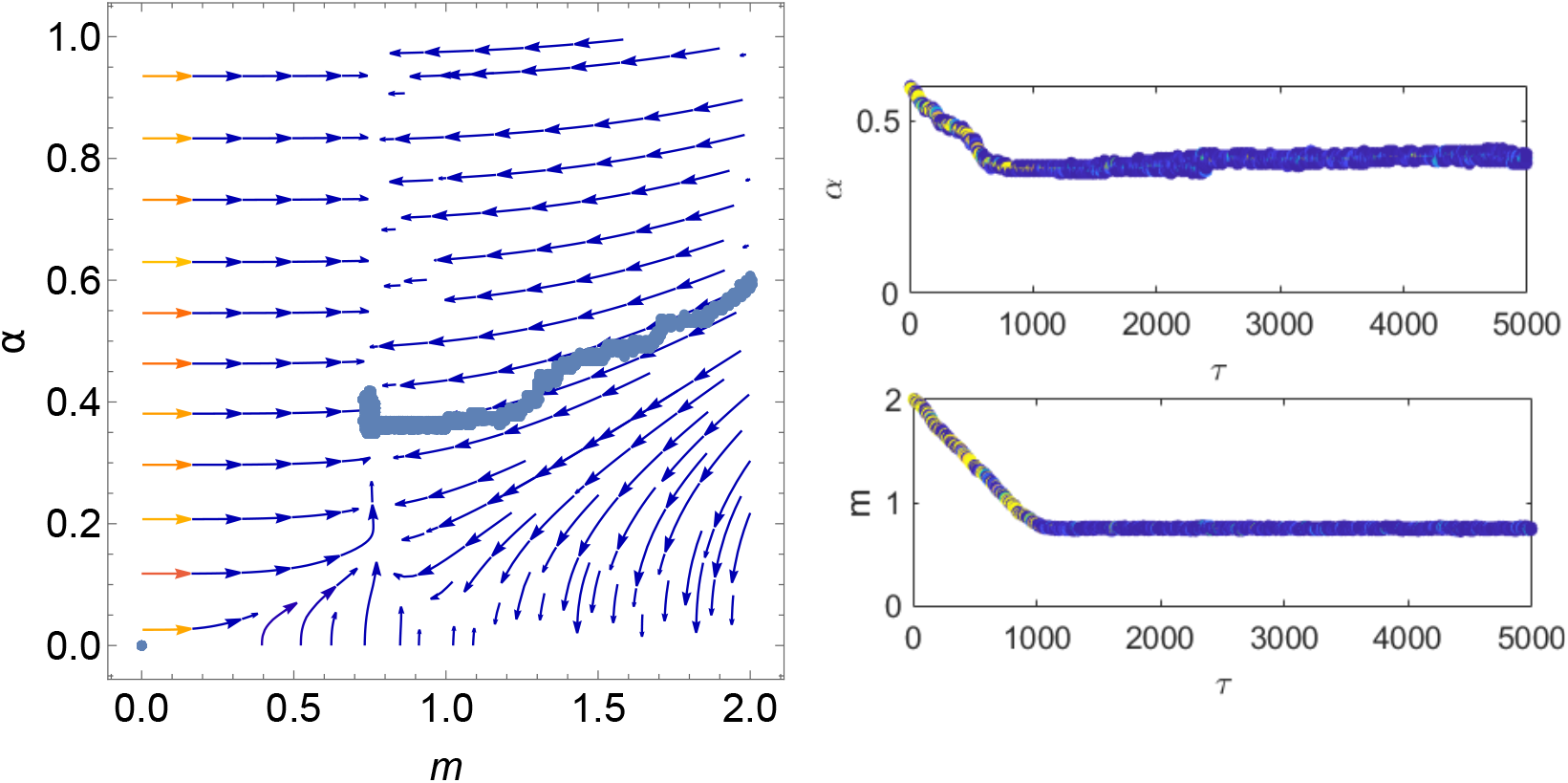
As in Figure D.15 but for Figure 4 (b).

**Figure D.18:**
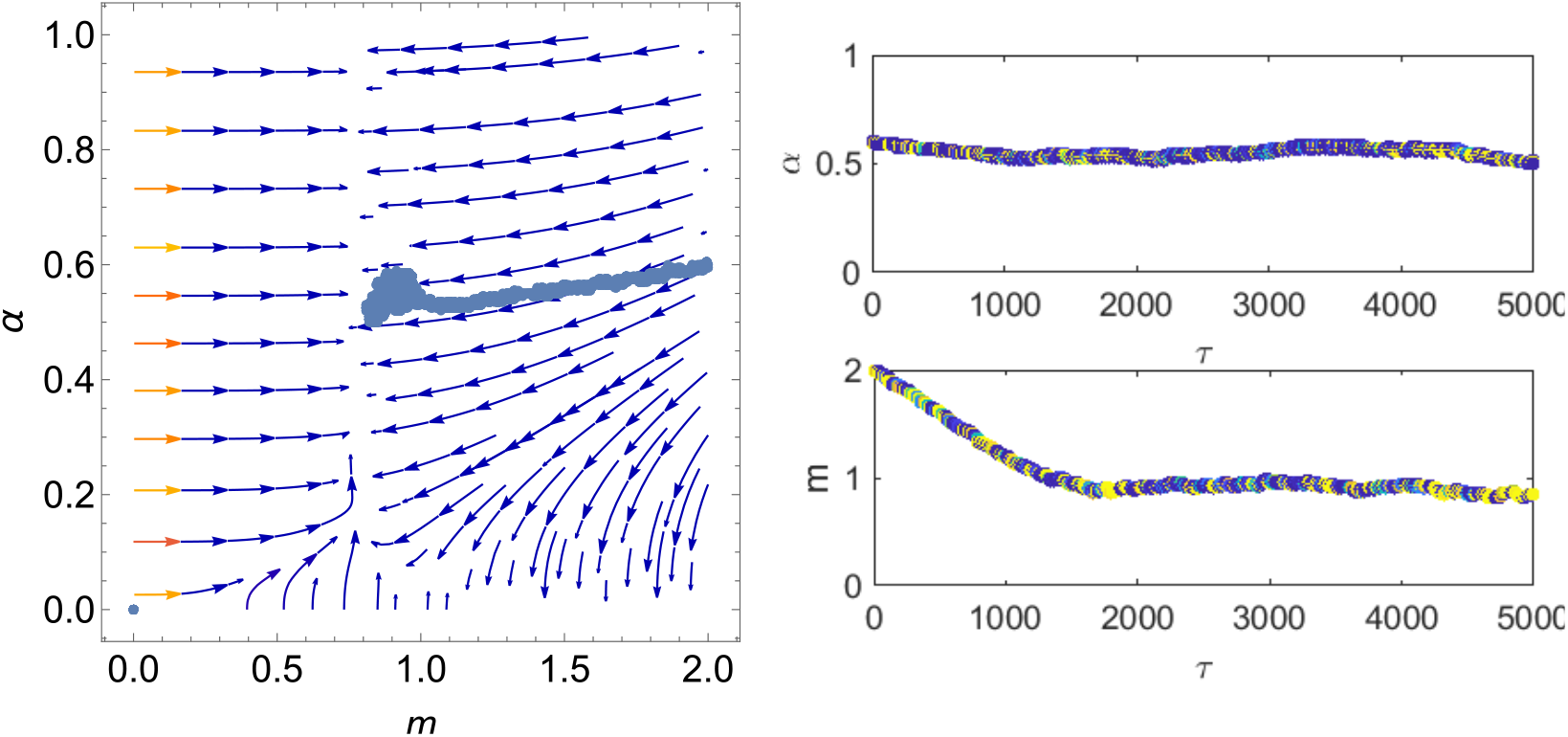
As in Figure D.16 but for Figure 4 (b).

**Figure D.19:**
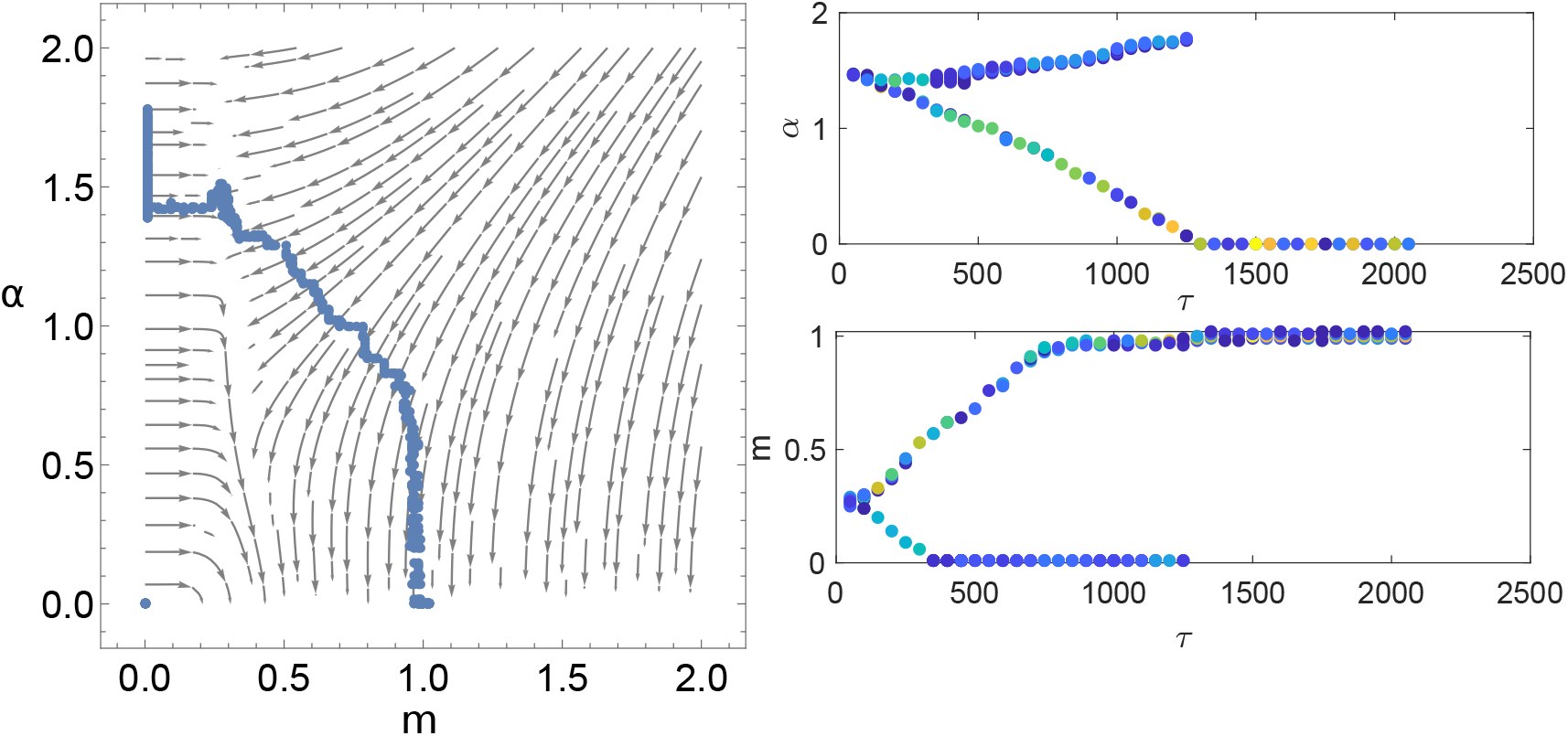
Numerical illustration of evolutionary branching for the case where *C* = 0.99. All other parameters the same as Figure 2, except (*m*(0), *α*(0)) = (0.25, 1.5), *δ* = 0.01, *f*_0_ = 0.05 and run for 2500/*μ* generations. Here, we observe stochastic extinction of the microgamete with large *α* due to their low frequency of approximately 0.01. Since branching is difficult to see using the *δ* and *f*_0_ in Figure 2, we used values that allow branching to be seen more easily.

**Figure D.20:**
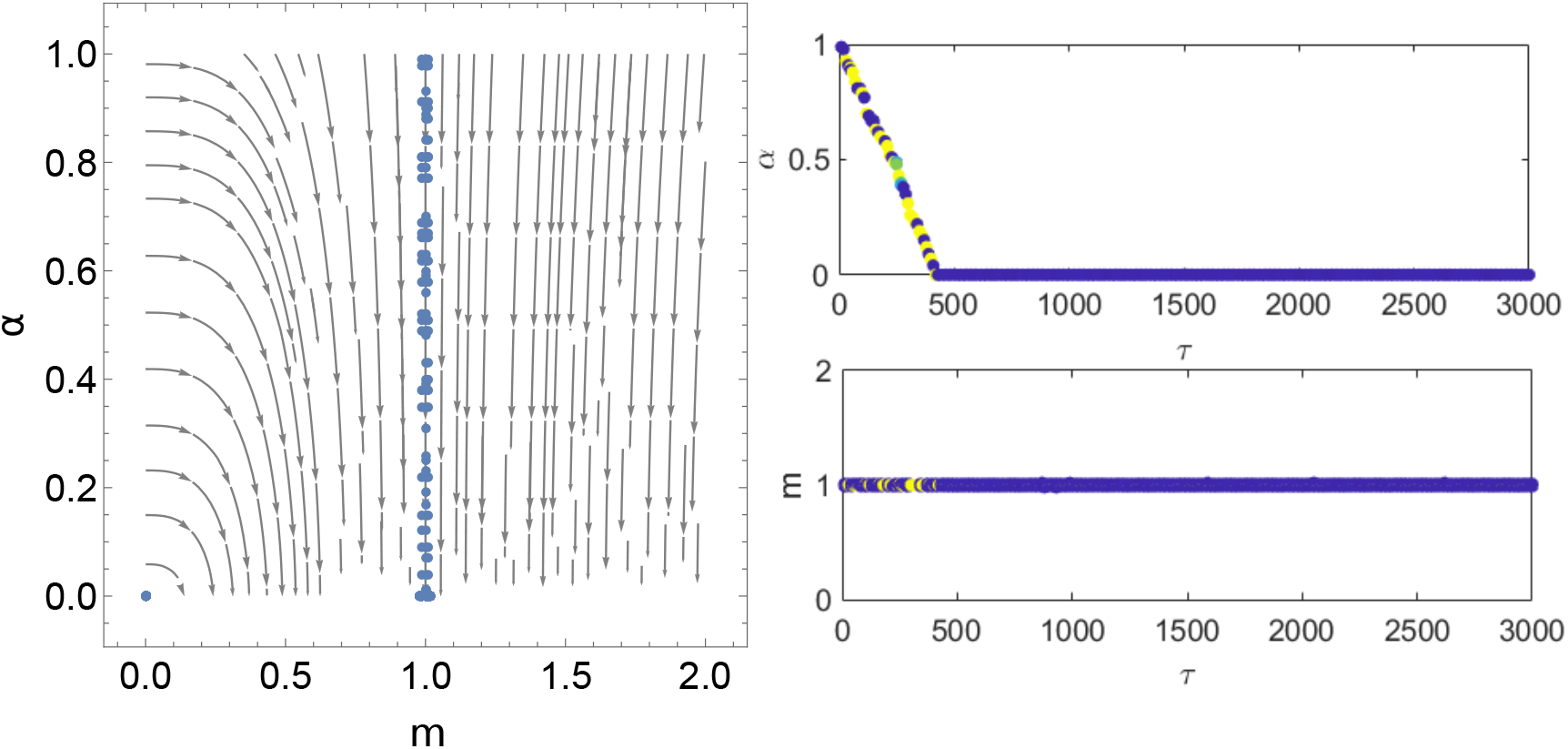
Numerical illustration of an absence of evolutionary branching for the case where *C* = 1. All other parameters the same as Figure 2, except (*m*(0), *α*(0)) = (1, 1), *δ* = 0.01, *f*_0_ = 0.05 and run for 3000/*μ* generations.

**Figure E.21:**
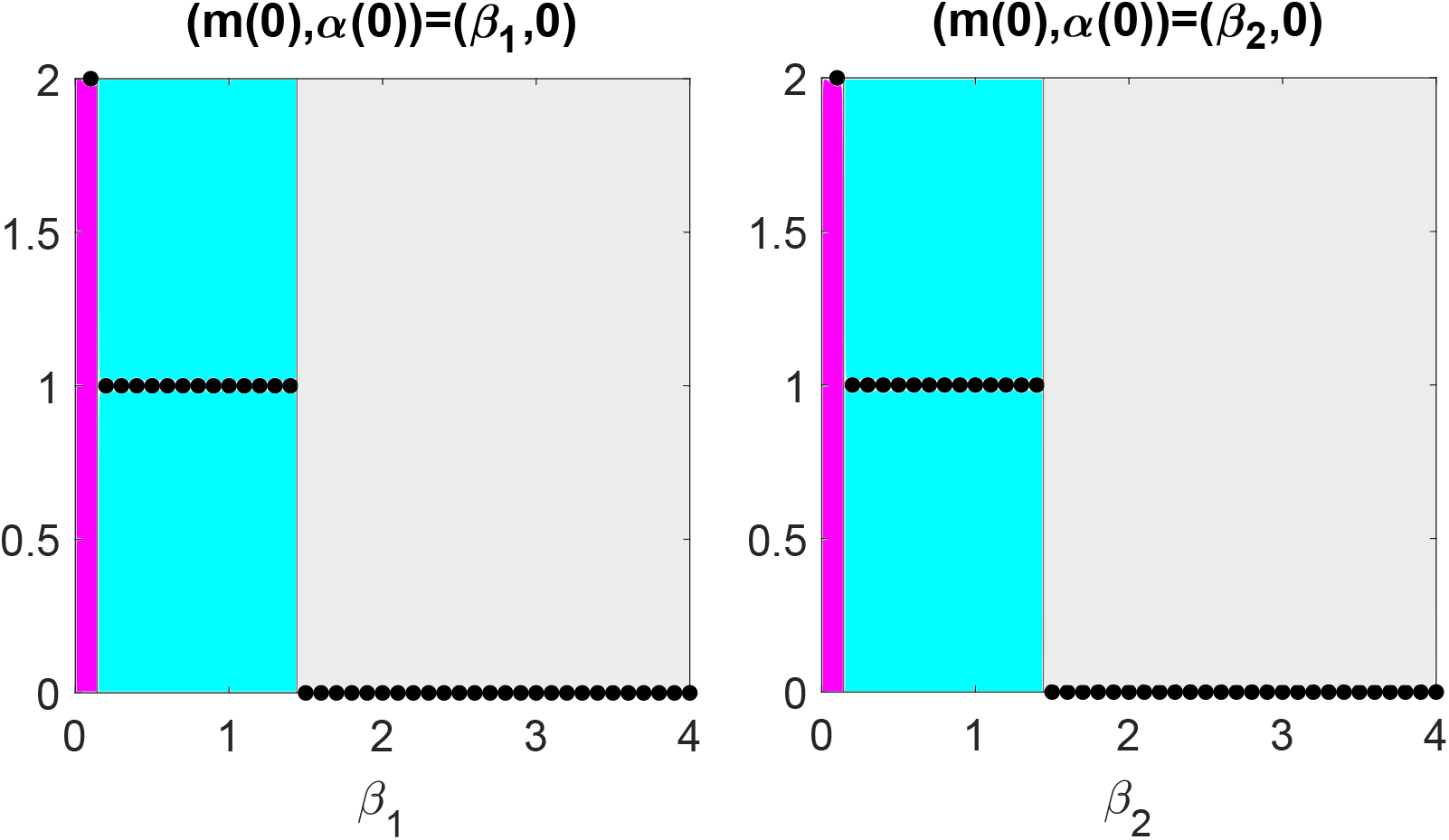
Black markers show numerical simulations of the evolutionary outcome in a switching environment with phenotypic plasticity. If the marker has a y-coordinate of 2, we evolve obligate fusion given both (*m*(0), *α*(0)) = (*β*_1_, 0) and (*m*(0), *α*(0)) = (*P*_1_*β*_1_ + (1 – *P*_1_)*β*_2_,0) for *i* ∈ {1, 2}. If the marker has a y-coordinate of 1, we evolve obligate fusion given (*m*(0), *α*(0)) = (*β*_1_, 0) only and if the marker sits at 0, we evolve no fusion given both initial conditions. Different coloured regions represent the behaviour predicted analytically predicted in Figure 6. In Panel (a) *β*_2_ = 2 and *i* = 1 and in Panel (b) *β*_1_ = 2 and *i* = 2. We see that the numerical simulations match the analytical predictions. The parameter conditions are *A* = 100, *M* = 1, *T* = 1, *C* = 0.5, *P*_1_ = 0.7 assuming *β*_1_ > *β*_2_, *δ* = 0.02, *μ* = 0.002 and *f*_0_ = 0.002. Simulation is run for 400/*μ* generations.

